# How Occam’s razor guides human decision-making

**DOI:** 10.1101/2023.01.10.523479

**Authors:** Eugenio Piasini, Shuze Liu, Pratik Chaudhari, Vijay Balasubramanian, Joshua I. Gold

**Affiliations:** International School for Advanced Studies (SISSA), Trieste, Italy; University of Pennsylvania, Philadelphia, PA, USA; PhD Program in Neuroscience, Harvard University, Boston, MA, USA; Santa Fe Institute, Santa Fe NM, USA; Rudolf Peierls Centre for Theoretical Physics, University of Oxford, Oxford, UK

## Abstract

Occam’s razor is the principle that, all else being equal, simpler explanations should be preferred over more complex ones. This principle is thought to guide human decision-making, but the nature of this guidance is not known. Here we used preregistered behavioral experiments to show that people tend to prefer the simpler of two alternative explanations for uncertain data. These preferences align quantitatively with predictions of formal theories of model selection that penalize excessive flexibility. These penalties emerge when considering not just the best explanation but the integral over all possible, relevant explanations. We further show that these simplicity preferences persist in humans, but not in certain artificial neural networks, even when they are maladaptive. Our results imply that principled notions of statistical model selection, including integrating over possible, latent causes to avoid overfitting to noisy data, may play a central role in human decision-making.

## Introduction

To make decisions in the real world, we must often choose between multiple, plausible explanations for noisy, sparse data. When evaluating competing explanations, Occam’s razor says that we should consider not just how well they account for data that have been observed, but also their potentially excessive flexibility in describing alternative, and potentially irrelevant, data that have not been observed (Baker, 2022) (e.g., “a ghost did it!”; Figure 1a,b). This kind of simplicity preference has long been proposed as an organizing principle for mental function (Feldman, 2016), such as in the early concept of Prägnanz in Gestalt psychology (Koffka, 2014; Wagemans et al., 2012), a number of “minimum principles” for vision (Hatfield, 1985) and theories that posit a central role for data compression in cognition (Chater, 1999; Chater & Vitányi, 2003). Simplicity has also been proposed as one of the key features that help us assess the merit of an explanation within broader theories of explanatory value (Wojtowicz & DeDeo, 2020).

**Figure 1.**
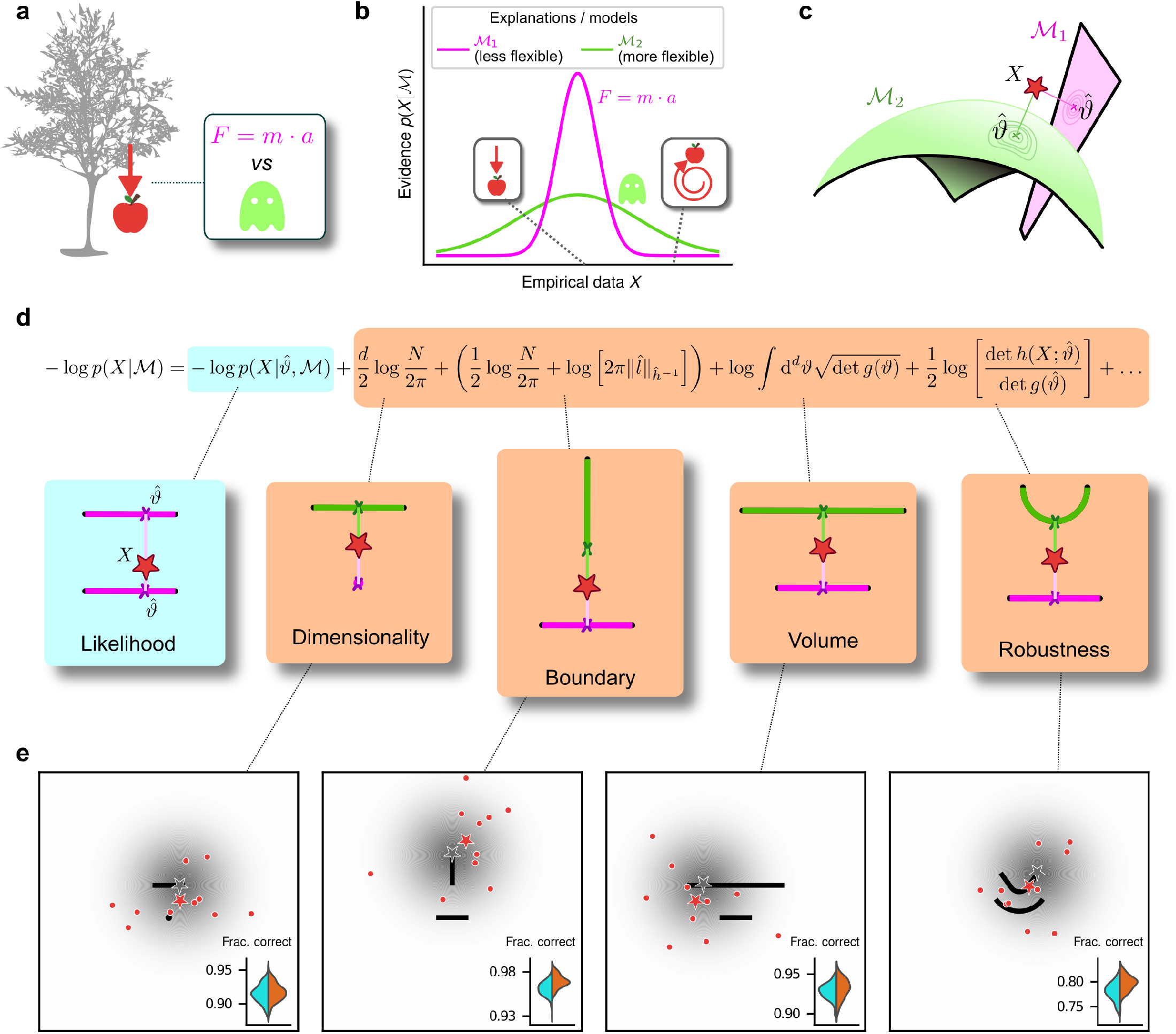
Formalizing Occam’s razor as Bayesian model selection to understand simplicity preferences in human decision-making. **a**: Occam’s razor prescribes an aversion to complex explanations (models). In Bayesian model selection, model complexity quantifies the flexibility of a model, or its capacity to account for a broad range of empirical observations. In this example, we observe an apple falling from a tree (left) and compare two possible explanations: 1) classical mechanics, and 2) the intervention of a ghost. **b**: Schematic comparison of the evidence of the two models in a. Classical mechanics (pink) explains a narrower range of observations than the ghost (green), which is a valid explanation for essentially any conceivable phenomenon (e.g., both a falling and spinning-upward trajectory, as in the insets). Absent further evidence and given equal prior probabilities, Occam’s razor posits that the simpler model (classical mechanics) is preferred, because its hypothesis space is more concentrated around the sparse, noisy data and thus avoids “overfitting” to noise. **c**: A geometrical view of the model-selection problem. Two alternative models are represented as geometrical manifolds, and the maximum-likelihood point for each model is represented as the projection of the data (red star) onto the manifolds. **d**: Systematic expansion of the log evidence of a model M (see previous work by Balasubramanian (1997) and Methods section A.2). 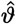 is the maximum-likelihood point on model ℳ for data X, N is the number of observations, d is the number of parameters of the model, 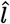 is the likelihood gradient evaluated at 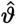, h is the observed Fisher information matrix, and g is the expected Fisher information matrix (see Methods). g(ϑ) captures how distinguishable elements of ℳ are in the neighborhood of ϑ (see Methods section A.2 and previous work by Balasubramanian (1997)). When M is the true source of the data X, h(X; ϑ) can be seen as a noisy version of g(ϑ), estimated from limited data (Balasubramanian, 1997). ĥ^−1^ is a shorthand for 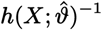, and 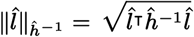 is the length of 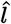 measured in the metric defined by ĥ^−1^. The ellipsis collects terms that decrease as N grows. Each term of the expansion represents a distinct geometrical feature of the model (Balasubramanian, 1997): dimensionality penalizes models with many parameters; boundary (a novel contribution of this work) penalizes models for which 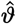 is on the boundary; volume counts the number of distinguishable probability distributions contained in 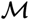; and robustness captures the shape (curvature) of ℳ near 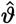 (see Methods section A.2 and previous work by Balasubramanian (1997)). **e**: Psychophysical task with variants designed to probe each geometrical feature in d. For each trial, a random location on one model was selected (gray star), and data (red dots) were sampled from a Gaussian centered around that point (gray shading). The red star represents the empirical centroid of the data, by analogy with c. The maximum-likelihood point can be found by projecting the empirical centroid onto one of the models. Participants saw the models (black lines) and data (red dots) only and were required to choose which model was best for the data. Insets: task performance for the given task variant, for a set of 100 simulated ideal Bayesian observers (orange) versus a set of 100 simulated maximum-likelihood observers (i.e., choosing based only on whichever model was the closest to the empirical centroid of the data on a given trial; cyan).

Numerous behavioral studies have shown that human decision-makers exhibit simplicity preferences when performing a variety of tasks including perceptual estimation, source reconstruction, and causal explanation (Chater, 1999; Genewein & Braun, 2014; Gershman & Niv, 2013; Griffiths & Tenenbaum, 2005; Johnson et al., 2014; Körding et al., 2007; Little & Shiffrin, 2009; Lombrozo, 2007; Pothos & Chater, 2002). These studies have proposed several formal definitions for what exactly may constitute the “simplicity” that is favored (or, equivalently, “complexity” that is disfavored) under these conditions, including the number of unobserved or unexplained causes invoked by an explanation (Pacer & Lombrozo, 2017), description length and Kolmogorov complexity (Chater, 1999), or statistical complexity arising from the computation of marginal likelihoods in Bayesian inference (Griffiths & Tenenbaum, 2005; Körding et al., 2007). Consistent with a Bayesian explanation, in some cases these kinds of simplicity preferences are diminished when more probabilistic evidence is provided, which has been proposed to reflect a Bayesian prior that places a heavier weight on simpler explanations (Bonawitz & Lombrozo, 2012; Lombrozo, 2007; Pacer & Lombrozo, 2017). Other human judgments have been shown to be compatible with a Bayesian framing that penalizes flexible explanations, where this flexibility is controlled by the size of its parameter space rather than its dimensionality (Blanchard et al., 2018). However, it remains unknown if and how these kinds of simplicity preferences relate quantitatively to the specific trade-offs between simplicity and goodness-of-fit of an explanation that are prescribed by theoretical notions of complexity.

The goal of the present study was to establish a theoretically grounded, quantitative account of simplicity biases in human perceptual decision-making. To do so, we devised a set of tasks that formalized these decisions as a statistical model-selection problem. This approach allowed us to identify specific features that make one model (i.e., a potential source of the observed data) more (or less) complex than another and thus require specific penalties to counteract their predicted effects on goodness-of-fit because of excessive flexibility (Balasubramanian, 1997). Critically, these penalties can depend on the data and therefore cannot be fully mimicked by a more general simplicity preference (Blanchard et al., 2018; Lombrozo, 2007). As we detail below, this approach allowed us to identify specific forms of simplicity biases that support the idea that Occam’s razor is not merely a preference but rather a manifestation of core decision processes that integrate over possible, latent causes to counteract the problems that arise when attempting to match flexible explanations to noisy data.

## Results

### Occam’s razor formalized as model selection

Given a set *X* of *N* observations and a set of possible parametric statistical models {ℳ_1_, ℳ_2_, …}, we seek the model ℳ that in some sense is the best description of the data *X*. In this context, Occam’s razor can be interpreted as penalizing how well a model fits the data by some measure of the model’s flexibility, or complexity, that could inappropriately inflate its goodness-of-fit, when comparing it against other models. Bayesian statistics provides such a measure of complexity and specifies how it should be traded off against goodness-of-fit to maximize decision accuracy. This trade-off occurs because although increased model complexity can provide a better match to more varied signals in the data, it also tends to cause errors by overfitting to noise (Balasubramanian, 1997; Good, 1968; Gull, 1988; Jaynes, 2003; H. Jeffreys, 1939; W. Jeffreys & Berger, 1991; MacKay, 1992; Smith & Spiegelhalter, 1980).

According to the Bayesian framework, models should be compared based on their evidence or marginal likelihood *p*(*X*|ℳ) = ∫*ϑw*(*ϑ*)*p*(*X*|ℳ, *ϑ*), where *ϑ* represents model parameters and *w*(*ϑ*) their associated prior. By varying the parameters, we can explore instances of a model and sweep out a manifold of possible descriptions of the data. Two such manifolds are visualized in Figure 1c, along with the maximum-likelihood parameters that assign the highest probability to the observed data. Under mild regularity assumptions and with sufficient data, the (log) evidence can be written as the sum of the maximum log likelihood of ℳ and several penalty factors (Figure 1d). These penalty factors, which are found even when the prior (data-independent) probabilities of the models under consideration are equal, can be interpreted as providing quantitatively defined preferences against certain models according to specific forms of complexity that they embody; i.e., a quantitative form of Occam’s razor (Balasubramanian, 1997; MacKay, 1992).

This approach, which we call the Fisher Information Approximation (FIA), has been used to identify worse-fitting, but better-generalizing, psychophysical models describing the relationship between physical variables (e.g., light intensity) and their psychological counterparts (e.g., brightness) (Myung et al., 2000). It is related to similar quantitative definitions of statistical model complexity, such as the Minimum Description Length (Grünwald, 2007; Lanterman, 2001; Rissanen, 1996), Minimum Message Length (Wallace, 2005), and Predictive Information (Bialek et al., 2001) frameworks. A key feature of the FIA is that if the prior over parameters *w*(*ϑ*) is taken to be uninformative (Jaynes, 2003), each penalty factor can be shown to capture a distinct geometric property of the model (Balasubramanian, 1997). Each of these geometric properties characterizes in a different way the degree to which the explanatory power of the model is concentrated nearby (or away from) the data.

Here we focus on the four leading terms (after the likelihood) of this Bayesian formulation, representing four distinct geometric properties of the model (Figure 1c,d). The first term is the model’s dimensionality (number of parameters), which is the well-known Bayesian Information Criterion (BIC) for model selection (Neath & Cavanaugh, 2012; Schwarz, 1978). The second term (a novel contribution of this work; see Methods section A.2) is a boundary penalty that measures the extent to which the set of possible observations that are well-explained by the model is bounded away from the data due to an abrupt cut-off in what the model can represent, which implies that the model concentrates its explanatory capacity in irrelevant regions of the space of possible observations. The third term is the model volume, which represents the number of distinguishable probability distributions it contains (Balasubramanian, 1997) and thus formalizes the idea of the “size of the parameter space” considered in previous studies Blanchard et al., 2018. The fourth term is related to the geometric shape of the model that is sensitive to whether the model tends to “bend away” from the data around the maximum-likelihood point. These last three terms (boundary, volume, and shape) all capture quantitatively the intuitive notion that among all the distributions described by the model family, only the ones that are closest to the data represent useful explanations, whereas the others constitute “wasted” explanatory power.

These complexity penalties emerge because the Bayesian framework marginalizes over the model parameters. In applying this framework to human decision-making, we interpret this marginalization as an integration over latent causes: to evaluate a particular explanation (or “model”) for a given set of observed data, one considers how likely the data are under that explanation, on average over all possible configurations of that explanation. Intuitively, flexible explanations are penalized by the averaging insofar as many of their configurations have nothing to do with the observed state of the world *X* and thus possess a vanishingly small likelihood *p*(*X* |ℳ, *ϑ*).

Consider the following example, in which the data are represented by a point on a plane, *X* = (*x, y*) (Figure 2, top left). The problem is to decide between two alternative explanations (models) for the data: 1) ℳ_1_, a Gaussian distribution centered in (0, 0) with unit, isotropic variance; and 2) ℳ_2_, a parametric family of Gaussians, also with unit variance, but with centers located anywhere along the straight line connecting (−1*/*2, 1) and (1*/*2, 1). It is clear that ℳ_2_ can explain a wider range of data, just like the ghost in Figure 1a, and is therefore more complex. For data that are equidistant from the two models, *X* = (0, 1*/*2), Occam’s razor prescribes that we should choose ℳ_1_. In other words, the decision boundary separating the area where one or the other model should be preferred is closer to ℳ_2_ (the more complex model) than ℳ_1_ (the simpler one). As illustrated by the simulations in Figure 2, this simplicity bias is specific to a decision-maker that integrates over the latent causes (model configurations) and does not result from sampling multiple possible explanations via other, less systematic means, for example by adding sensory and/or choice noise (see also Supplementary Figure B.9).

**Figure 2.**
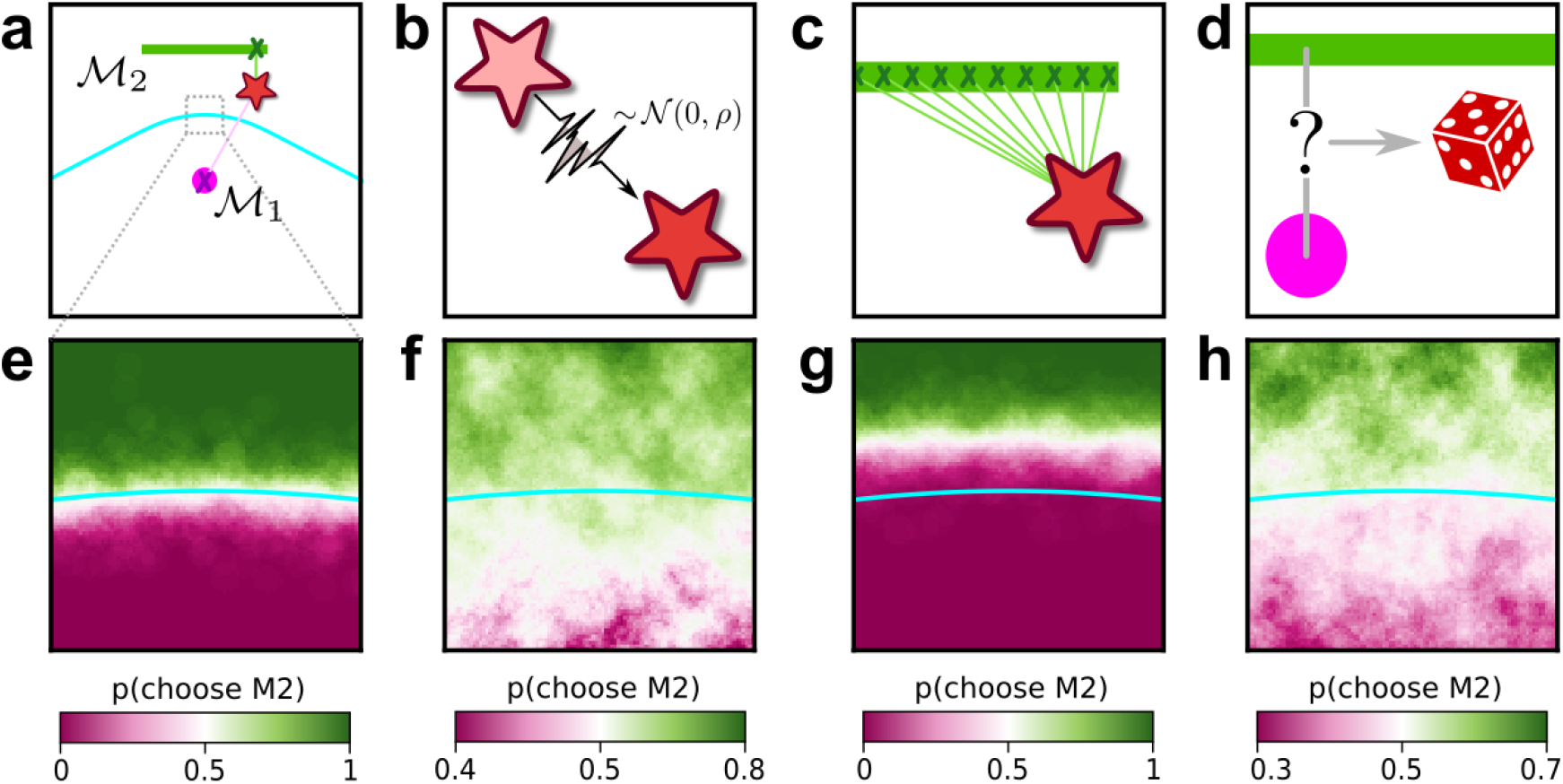
Integration over latent causes leads to Occam’s razor. **a**: Schematic of a simple decision-making scenario. A single datapoint (star) is sampled from one of two models (pink dot, green bar). One of the models (ℳ_1_) is a Gaussian with known variance, centered at the location of the pink dot. The other model (ℳ_2_) is a parametric family of Gaussians, with known and fixed variance and center located at a latent location along the green bar. Cyan line: boundary indicating locations in data space that are equidistant from ℳ_1_ and ℳ_2_. **b-d**: Potential components of a decision-making observer for this scenario, which we call Noise-Integration-Noise observer (see Methods section A.1 and Supplementary Information section B.7 for further details). **b**: Sensory noise: the observer does not have access to the true data (location of the star), but a noisy version of it corrupted by Gaussian noise with variance ρ. **c**: Integration over latent causes: the observer can consider possible positions of the center of the Gaussian in model ℳ_2_. **d**: Choice noise: after forming an internal estimate of the relative likelihood of ℳ_1_ and ℳ_2_, the observer can choose a model based on a deterministic process (for instance, always pick the most likely one), or a stochastic one where the probability of sampling one model is related to its likelihood. **e-h**: Behavior of the observer as a function of the location of the datapoint, within the zoomed-in region highlighted in a, and of the presence of the mechanisms illustrated in b-d. **e**: probability that the observer will report ℳ_2_ as a function of the location of the datapoint, when sensory and choice noise are low and in absence of integration over latent causes. **f**: same as e, but with strong sensory noise. The decision boundary of the observer (white area) is shifted towards the simplest model (ℳ _1_) compared to e. This shift means that, when the data is equidistant from ℳ_1_ and ℳ_2_, the observer prefers the more complex model (ℳ _2_). **g**: same as e, but in presence of integration over latent causes. The decision boundary of the observer is shifted in the opposite direction as f, and when the data is equidistant from both models the observer prefers the simplest model. **h**: same as e, but with strong choice noise. Choice noise has no effect on the location of the decision boundary.

### Humans exhibit theoretically grounded simplicity preferences

To relate the FIA complexity terms to the potential preferences exhibited by both human and artificial decision-makers, we designed a simple decision-making task. For each trial, *N* = 10 simultaneously presented, noisy observations (red dots in Figure 1e) were sampled from a 2D Normal (“generative”) distribution centered somewhere within one of two possible shapes (black shapes in Figure 1e). The identity of the shape generating the data (top versus bottom) was chosen at random with equal probability. Likewise, the location of the center of the Normal distribution within the selected shape was sampled uniformly at random, in a way that did not depend on the model parameterization, by using Jeffreys prior (Jaynes, 2003). Given the observations, the decision-maker chose the shape (model) that was more likely to contain the center of the generative distribution.

We used four task variants, each designed to probe primarily one of the distinct geometrical features that are penalized in Bayesian model selection. In other words, for each variant, a Bayesian observer is expected to have a particular, quantitative preference away from the more-complex alternative in each pair (Figure 1c and d). For this task, the FIA provided a good approximation of the exact Bayesian posterior (Supplementary Information section B.1). Accordingly, simulated observers that increasingly integrated over latent causes, like the Bayesian observer, exhibited increasing FIA-like biases. These biases were distinguishable from (and degraded by) effects of increasing sensory and/or choice noise (Supplementary Information B.7.1 and Supplementary Figure B.10).

For our human studies, we used the on-line research platform Pavlovia to implement the task, and Prolific to recruit and test participants. Following our preregistered approaches (Piasini et al., 2020, 2021), we collected data from 202 participants, divided into four groups that each performed one of the four separate versions of the task depicted in Figure 1e (each group comprised ~50 participants). We provided instructions that used the analogy of seeds from a flower located in one of two flower beds, to provide an intuitive framing of the key concepts of noisy data generated by a particular instance of a parametric model from one of two model families. To minimize the possibility that participants would simply learn from implicit or explicit feedback over the course of each session to make more optimal (i.e., simplicity-preferring) choices of flower beds, we: ∼1) used conditions for which the difference in performance between ideal observers that penalized model complexity according to the FIA and simulated observers that used only model likelihood was ∼1% (depending on the task type; Figure 1e, insets), which translates to ∼5 additional correct trials over the course of an entire experiment; and 2) provided feedback only at the end of each block of 100 trials, not each trial.

Simple behavioral summaries show that for ambiguous trials, in which the likelihood of either model was similar (namely, where the data was equidistant from both models, corresponding to the “all else being equal” condition of the classical statement of Occam’s razor), participants chose more often the simplest alternative (pink area extending above the cyan curves in Figure 3a). To test this tendency across all stimulus conditions, including when one model had a higher likelihood than the other, and to assess quantitatively the degree to which each participant’s choices were affected by model likelihood and each of the FIA features, we analyzed our data using a hierarchical (Bayesian) logistic regression approach (see Methods section A.6). We defined each participant’s sensitivity to each FIA term as a normalized quantity, relative to their likelihood sensitivity (i.e., by dividing the logistic coefficient associated with a given FIA term by the logistic coefficient associated with the likelihood). This analysis showed that the participants were sensitive to all four forms of model complexity (Figure 3). Specifically, the estimated normalized population-level sensitivity (posterior mean ± st. dev., where zero implies no sensitivity and one implies Bayes-optimal sensitivity) was 4.66±0.96 for dimensionality, 1.12±0.10 for boundary, 0.23±0.12 for volume, and 2.21±0.12 for robustness (note that, following our preregistered plan, we emphasize parameter estimation using Bayesian approaches (Gelman et al., 2014; Kruschke, 2015; McElreath, 2016) here and throughout the main text, and we provide complementary null hypothesis significance testing in the Supplementary Information section B.6 and Table B.8).

**Figure 3.**
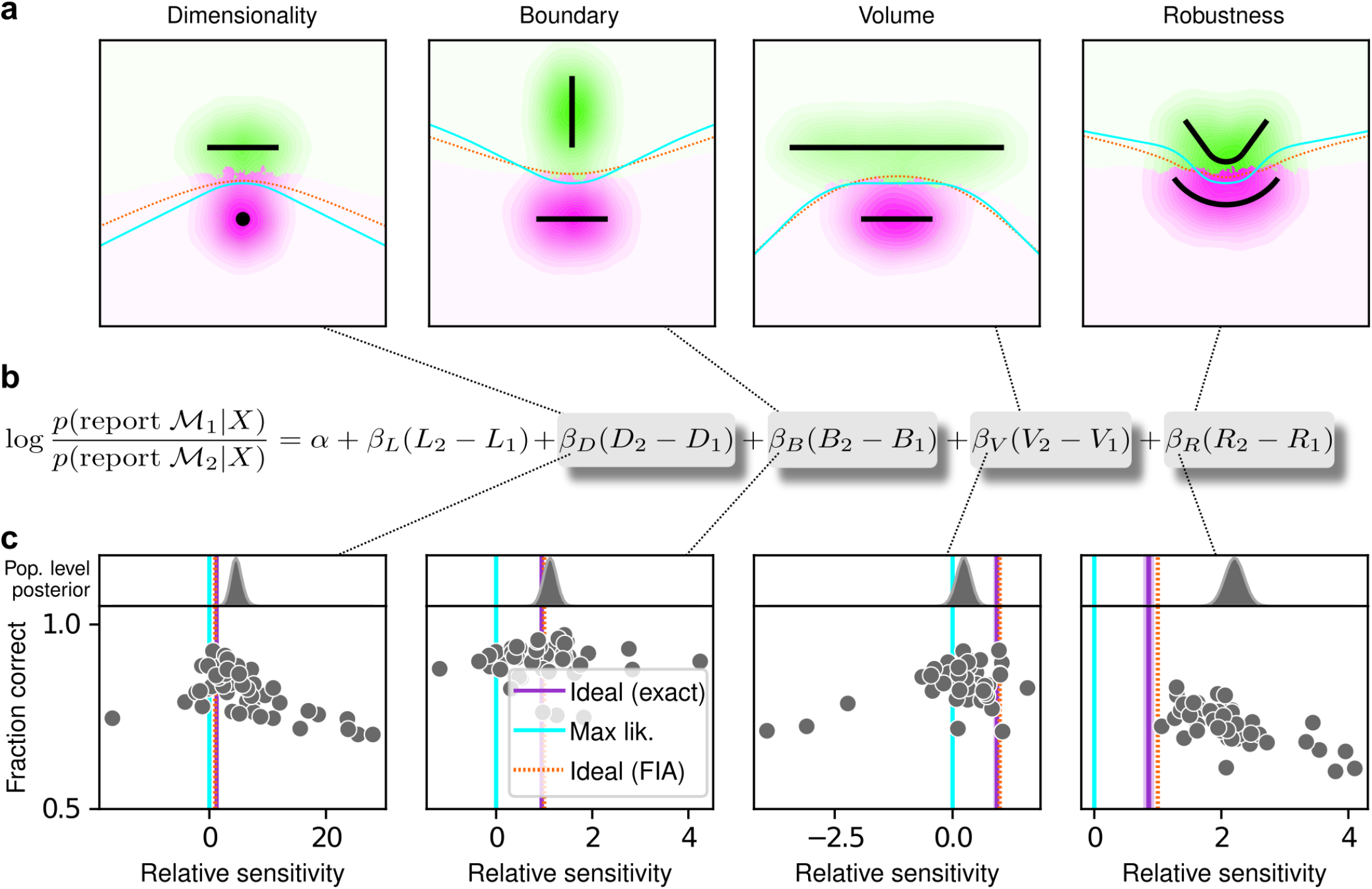
Humans exhibit theoretically grounded simplicity preferences. **a**: Summary of human behavior. Hue (pink/green): k-nearest-neighbor interpolation of the model choice, as a function of the empirical centroid of the data. Color gradient (light/dark): marginal density of empirical data centroids for the given model pair, showing the region of space where data were more likely to fall. Cyan solid line: decision boundary for an observer that always chooses the model with highest maximum likelihood. Orange dashed line: decision boundary for an ideal Bayesian observer. The participants’ choices tended to reflect a preference for the simpler model, particularly near the center of the screen, where the evidence for the alternatives was weak. For instance, in the left panel there is a region where data were closer to the line than to the dot, but participants chose the dot (the simpler, lower-dimensional “model”) more often than the line. **b**: Participant sensitivity to each geometrical feature characterizing model complexity was estimated via hierarchical logistic regression (see Methods section A.6 and Supplementary Information section B.2), using as predictors a constant to account for an up/down choice bias, the difference in likelihoods for the two models (L_2_ − L_1_) and the difference in each FIA term for the two models (D_2_ − D_1_, etc). Following a hierarchical regression scheme, the participant-level sensitivities were in turn modeled as being sampled from a population-level distribution. The mean of this distribution is our population-level estimate for the sensitivity. **c**: Overall accuracy versus estimated relative FIA sensitivity for each task condition, as indicated. Points are data from individual participants. Each fitted FIA coefficient was normalized to the likelihood coefficient and thus could be interpreted as a relative sensitivity to the associated FIA term. For each term, an ideal Bayesian observer would have a relative sensitivity of one (dashed orange lines), whereas an observer that relied on only maximum-likelihood estimation (i.e., choosing “up” or “down” based on only the model that was the closest to the data) would have a relative sensitivity of zero (solid cyan lines). Top, gray: Population-level estimates (posterior distribution of population-level relative sensitivity given the experimental observations). Bottom: each gray dot represents the task accuracy of one participant (y axis) versus the posterior mean estimate of the relative sensitivity for that participant (x axis). Intuitively, the population-level posterior can be interpreted as an estimate of the location of the center of the cloud of dots representing individual subjects in the panel below. See Methods section A.6 for further details on statistical inference and the relationship between population-level and participant-level estimates. Purple: relative sensitivity of an ideal observer that samples from the exact Bayesian posterior (not the approximated one provided by the FIA). Shading: posterior mean ± 1 or 2 stdev., estimated by simulating 50 such observers.

We complemented these analyses with formal model comparisons (WAIC; see Supplementary Information section B.6.1 and Tables B.6 and B.7), which confirmed that their behavior was better described by taking into account the geometric penalties defined by the theory of Bayesian model selection, rather than by relying only on the minimum distance between model and data (i.e., the maximum-likelihood solution). Their decisions also were consistent with processes that tended to integrate over latent causes (and tended to exhibit moderate levels of sensory noise and low choice noise; Supplementary Information B.7 and Supplementary Figures B.11 and B.12). Overall, our analyses indicate that people tend to integrate over latent causes in a way that gives rise to Occam’s razor, manifesting as sensitivity to the geometrical features in Bayesian model selection that characterize model complexity.

The participants also exhibited substantial individual variability in performance that included ranges of sensitivities to each FIA term that spanned optimal and sub-optimal values. This variability was large compared to the uncertainty associated with participant-level sensitivity estimates (Supplementary Information B.4) and impacted performance in a manner that highlighted the usefulness of appropriately tuned (i.e., close to Bayes optimal) simplicity preferences: accuracy tended to decline for participants with FIA sensitivities further away from the theoretical predictions (Figure 2c; posterior mean ± st. dev. of Spearman’s rho between accuracy and |*β* − 1|, where *β* is the sensitivity: dimensionality, -0.69±0.05; boundary, -0.21±0.11; volume, -0.10±0.10; robustness, -0.54±0.10). The sub-optimal sensitivities exhibited by many participants did not appear to result simply from a lack of task engagement, because FIA sensitivity did not correlate with errors on easy trials (posterior mean ± st. dev. of Spearman’s rho between lapse rate, estimated with an extended regression model detailed in Methods section A.6.1, and the absolute difference from optimal sensitivity for: dimensionality, 0.08±0.12; boundary, 0.15±0.12; volume, -0.04±0.13; robustness, 0.15±0.14; see Supplementary Information section B.5). Likewise, sub-optimal FIA sensitivity did not correlate with weaker likelihood sensitivity for the boundary (rho=-0.13±0.11) and volume (-0.06±0.11) terms, although stronger, negative relationships with the dimensionality (-0.35±0.07) and robustness terms (-0.56±0.10) suggest that the more extreme and variable simplicity preferences under those conditions (and lower performance, on average; see Figure 2c) reflected a more general difficulty in performing those versions of the task.

### Artificial Neural Networks learn optimal simplicity preferences

To better understand the optimality, variability, and generality of the simplicity preferences exhibited by our human participants, we compared their performance to that of artificial neural networks (ANNs) trained to optimize performance on this task. We used a novel ANN architecture that we designed to perform statistical model selection, in a form applicable to the task described above (Figure 4a,b). On each trial, the network took as input two images representing the models to be compared, and a set of coordinates representing the observations on that trial. The output of the network was a decision between the two models, encoded as a softmax vector. We analyzed 50 instances of the ANN that differed only in the random initialization of their weights and in the examples seen during training, using the same logistic-regression approach we used for the human participants.

**Figure 4.**
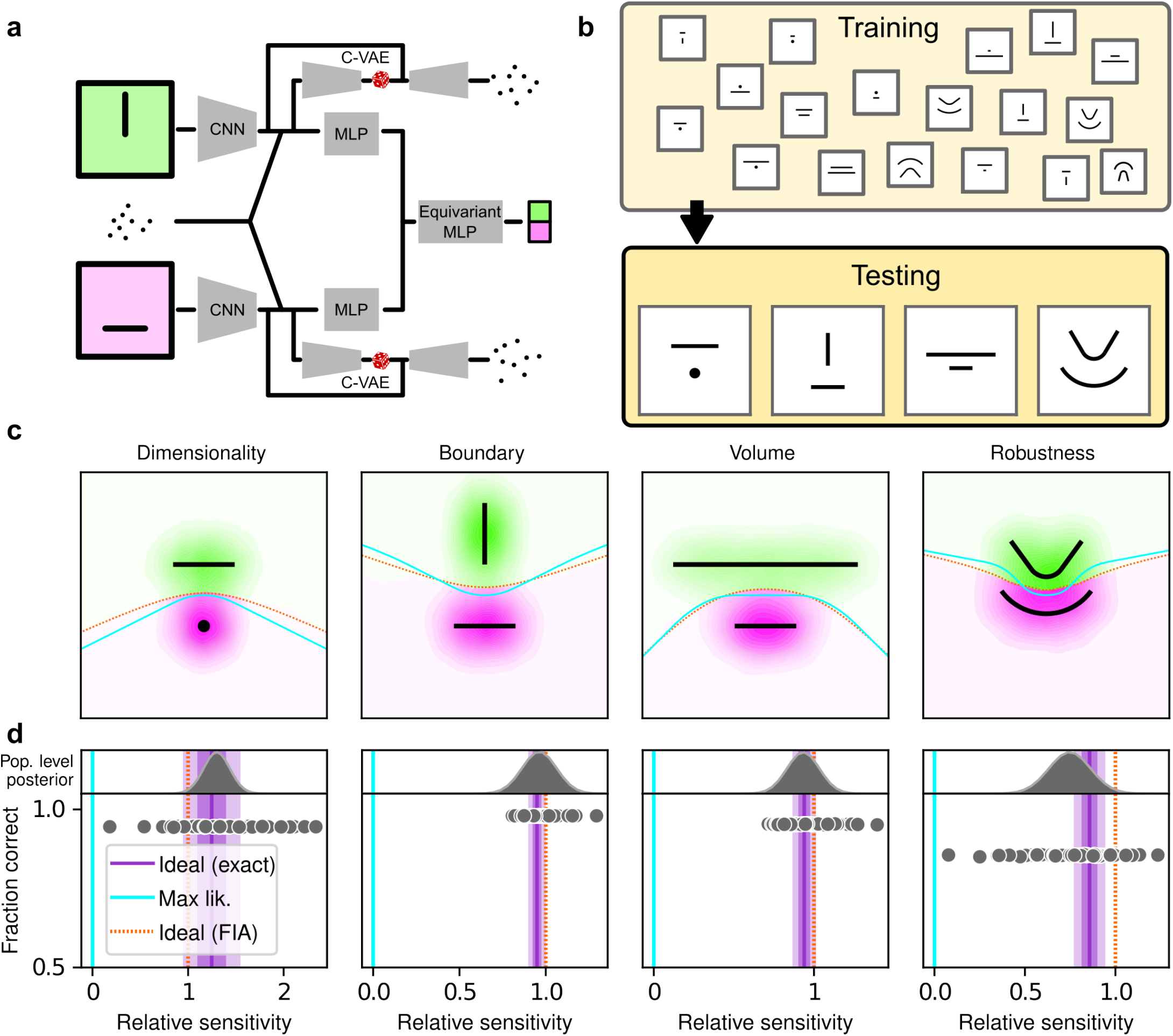
Artificial neural networks exhibit theoretically grounded simplicity preferences. **a**: A novel deep neural-network architecture for statistical model selection. The network (see text and Methods for details) takes two images as input, each representing a model, and a set of 2D coordinates, each representing a datapoint. The output is a softmax-encoded choice between the two models. **b**: Each network was trained on multiple variants of the task, including systematically varied model length or curvature, then tested using the same configurations as for the human studies. **c**: Summary of network behavior, like Figure 3a. Hue (pink/green): k-nearest-neighbor interpolation of the model choice, as a function of the empirical centroid of the data. Color gradient (light/dark): marginal density of empirical data centroids for the given model pair, showing the region of space where data were more likely to fall. Cyan solid line: decision boundary for an observer that always chooses the model with highest maximum likelihood. Orange dashed line: decision boundary for an ideal Bayesian observer. **d**: Estimated relative sensitivity to geometrical features characterizing model complexity. As for the human participants, each fitted FIA coefficient was normalized to the likelihood coefficient and thus can be interpreted as a relative sensitivity to the associated FIA term. For each term, an ideal Bayesian observer would have a relative sensitivity of one (dashed orange lines), whereas an observer that relied on only maximum-likelihood estimation (i.e., choosing”up” or “down” based on only the model that was the closest to the data) would have a relative sensitivity of zero (solid cyan lines). Top: population-level estimate (posterior distribution of population-level relative sensitivity given the experimental observations; see Methods section A.6 for details). Bottom: each gray dot represents the task accuracy of one of 50 trained networks (y axis) versus the posterior mean estimate of the relative sensitivity for that network (x axis). Intuitively, the population-level posterior can be interpreted as an estimate of the location of the center of the cloud of dots representing individual subjects in the panel below. See Methods section A.6 for further details on statistical inference and the relationship between population-level and participant-level estimates. Purple: relative sensitivity of an ideal observer that samples from the exact Bayesian posterior (not the approximated one provided by the FIA). Shading: posterior mean ± 1 or 2 stdev., estimated by simulating 50 such observers.

The ANN was designed as follows (see Methods for more details). The input stage consisted of two pretrained VGG16 convolutional neural networks (CNNs), each of which took in a pictorial representation of one of the two models under consideration. VGG was chosen as a popular architecture that is often taken as a benchmark for comparisons with the human visual system (Muratore et al., 2022; Schrimpf et al., 2020). The CNNs were composed of a stack of convolutional layers whose weights were kept frozen at their pretrained values, followed by three fully-connected layers whose weights were allowed to change during training. The outputs of the CNNs were each fed into a multilayer perceptron (MLP) consisting of linear, rectified-linear (ReLU), and batch-normalization layers. The MLP outputs were then concatenated and fed into an equivariant MLP, which enforces equivariance of the network output under position swap of the two models through a custom parameter-sharing scheme (Ravanbakhsh et al., 2017). The network also contained two conditional variational autoencoder (C-VAE) structures, which sought to replicate the data-generation process conditioned on each model and therefore encouraged the fully connected layers upstream to learn model representations that captured task-relevant features.

After training, the ANNs performed the task substantially better than the human participants, with higher overall accuracies that included higher likelihood sensitivities (Supplementary Information section B.3) and simplicity preferences that more closely matched the theoretically optimal values (Figure 4c,d). These simplicity preferences were closer to those expected from simulated observers that use the exact Bayesian model posterior rather than the FIA-approximated one, consistent with the fact that the FIA provides an imperfect approximation to the exact Bayesian posterior. These simplicity preferences varied slightly in magnitude across the different networks, but unlike for the human participants this variability was relatively small (compare ranges of values in Figures 3c and 4d, plotted on different *x*-axis scales) and it was not an indication of suboptimal network behavior because it was not related systematically to any differences in the generally high accuracy rates for each condition (Figure 4d; posterior mean ± st. dev. of Spearman’s rho between accuracy and |*β* − 1|, where *β* is the sensitivity: dimensionality, -0.14±0.10; boundary, 0.08±0.11; volume, -0.12±0.11; robustness, -0.08±0.11). These results imply that the stochastic nature of the task gives rise to some variability in simplicity biases even after extensive training to optimize performance accuracy, but this source of variability cannot by itself account for the range of sensitivities (and suboptimalities) exhibited by the human participants.

### Humans, unlike ANNs, maintain suboptimal simplicity preferences

Our results, combined with the fact that we did not provide trial-by-trial feedback to the participants while they performed the task, suggest that the human simplicity preferences we measured were not simply learned optimizations for these particular task conditions but rather are a more inherent (and variable) part of how we make decisions under uncertainty. However, because we provided each participant with instructions that echoed Bayesian-like reasoning (see Methods) and a brief training set with feedback before their testing session, we cannot rule out from the data presented in Figure 3 alone that at least some aspects of the simplicity preferences we measured from the human participants depended on those specific instructions and training conditions. We therefore ran a second experiment to rule out this possibility. For this experiment, we used the same task variants as above but a different set of instructions and training, designed to encourage participants to pick the model with the maximum likelihood (i.e., not integrate over latent causes but instead just consider the single cause that best matches the observed data), thus disregarding model complexity. Specifically, the visual cues were the same as in the original experiment, but the participants were asked to report which of the two shapes on the screen was closest to the center-of-mass of the dot cloud. We ensured that the participants recruited for this “maximum-likelihood” task had not participated in the original, “generative” task. We again followed a preregistered plan for the experiment and analysis (Piasini et al., 2022), and we collected data from 201 participants, divided in four groups of ~50 participants each. We also trained and tested ANNs on this version of the task, using the maximum-likelihood solution as the correct answer.

Despite this major difference in instructions and training, the human participants exhibited similar simplicity preferences on the generative and maximum-likelihood tasks, suggesting that humans have a general predilection for simplicity even without relevant instructions or incentives (Figure 5, left). Specifically, despite some quantitative differences, the distributions of relative sensitivities showed the same basic patterns for both tasks, with a general increase of relative sensitivity from volume (0.19±0.08 for the maximum-likelihood task; compare to values above), to boundary (0.89±0.10), to robustness (2.27±0.15), to dimensionality (2.29±0.41). In stark contrast to the human data and to ANNs trained on the true generative task, ANN sensitivity to model complexity on the maximum-likelihood task was close to zero for all four terms (Figure 5, right).

**Figure 5.**
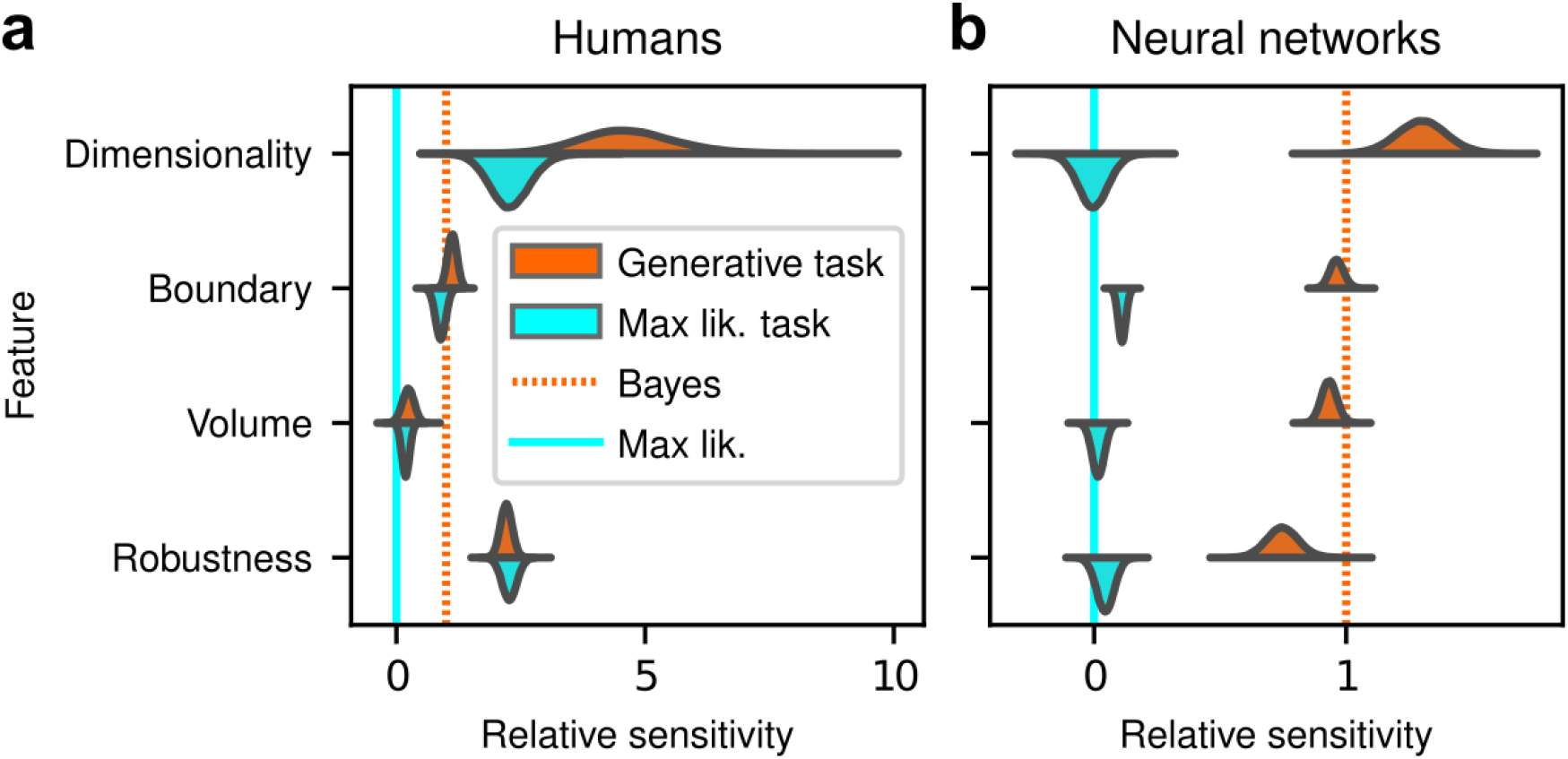
Humans, but not artificial networks, exhibit simplicity preferences even when they are suboptimal. **a**: Relative sensitivity of human participants to the geometric complexity terms (population-level estimates, as in Figure 3c, top) for two task conditions: 1) the original, “generative” task where participants were implicitly instructed to solve a model-selection problem (same data as in Figure 3c, top; orange); and 2) a “maximum-likelihood” task variant, where participants were instructed to report which of two models has the highest likelihood (shortest distance from the data; cyan). The two task variants were tested on distinct participant pools of roughly the same size (202 participants for the generative task, 201 for the maximum-likelihood task, in both cases divided in four groups of roughly 50 participants each). Solid cyan lines: relative sensitivity of a maximum-likelihood observer. Orange dashed lines: relative sensitivity of an ideal Bayesian observer. **b**: Same comparison and format (note the different x-axis scaling), but for two distinct populations of 50 deep neural networks trained on the two variants of the task (orange is the same data as in Figure 4d, top).

To summarize the similarities and differences between how humans and ANNs used simplicity biases to guide their decision-making behaviors for these tasks, and their implications for performance, Figure 6 shows overall accuracy for each set of conditions we tested. Specifically, for each participant or ANN, task configuration, and instruction set, we computed the percentage of correct responses with respect to both the generative task (i.e., for which theoretically optimal performance depends on simplicity biases) and the maximum-likelihood task (i.e., for which theoretically optimal performance does not depend on simplicity biases). Because the maximum-likelihood solutions are deterministic (they depend only on which model the data centroid is closest to, and thus there exists an optimal, sharp decision boundary that leads to perfect performance) and the generative solutions are not (they depend probabilistically on the likelihood and bias terms, so it is generally impossible to achieve perfect performance), performance on the former is expected to be higher than on the latter. Accordingly, both ANNs and (to a lesser extent) humans tended to perform better when assessed relative to maximum-likelihood solutions. Moreover, the ANNs tended to exhibit behavior that was consistent with optimization to the given task conditions: networks trained to find maximum-likelihood solutions did better than networks trained to find generative solutions for the maximum-likelihood task, and networks trained to find generative solutions did better than networks trained to find maximum-likelihood solutions for the generative task. In contrast, humans tended to adopt similar strategies regardless of the task conditions, in all cases using Bayesian-like simplicity biases.

**Figure 6.**
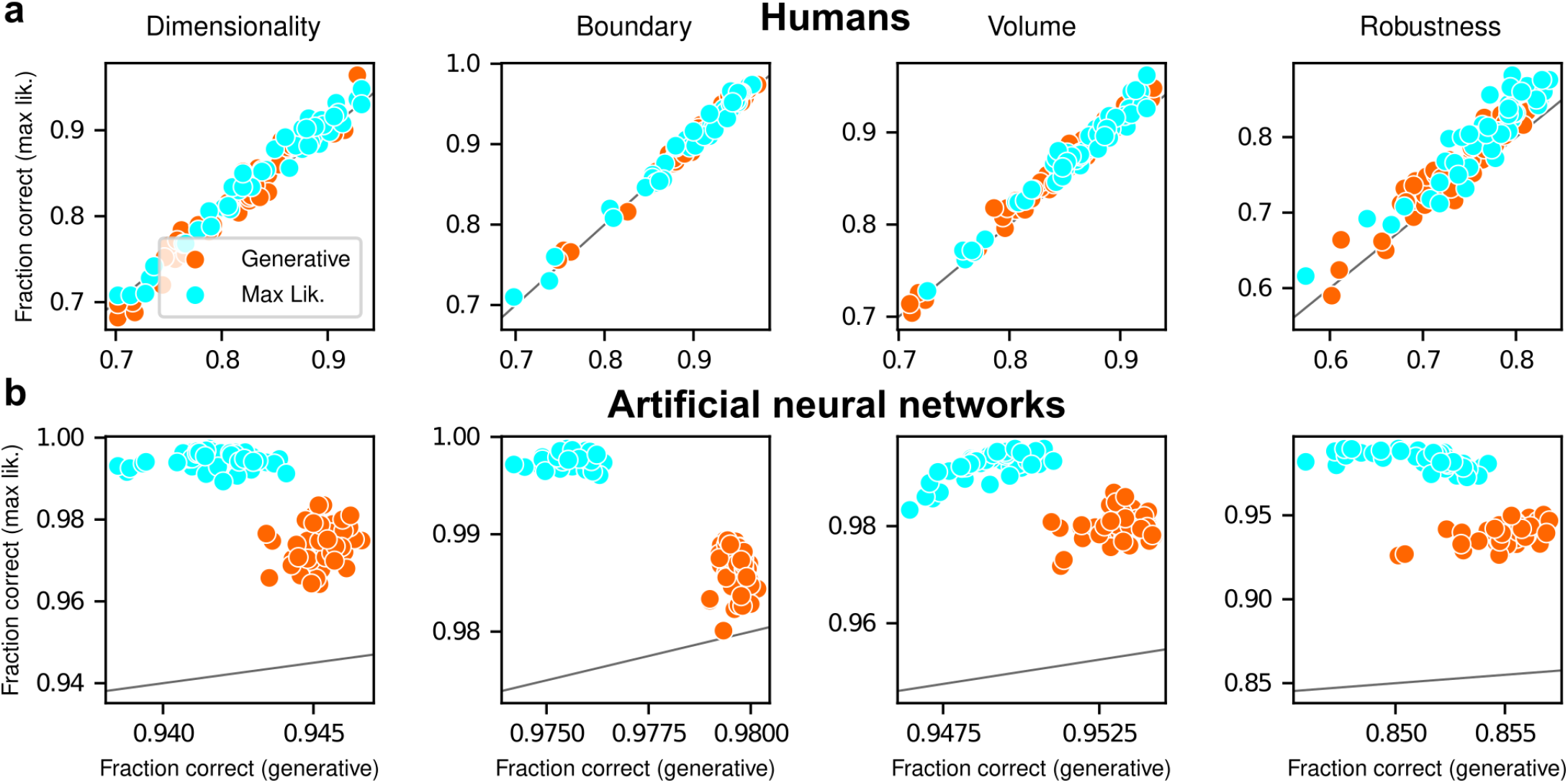
Humans and artificial neural networks have different patterns of accuracy reflecting their different use of simplicity preferences. Each panel shows accuracy with respect to maximum-likelihood solutions (i.e., the model closest to the centroid of the data; ordinate) versus with respect to generative solutions (i.e., the model that generated the data; abscissa). The gray line is the identity. Columns correspond to the four task variants associated with the four geometric complexity terms, as indicated. **a**: Data from individual human participants (points), instructed to find the generative (orange) or maximum-likelihood (cyan) solution. Human performance was higher when evaluated against maximum-likelihood solutions than it was when evaluated against generative solutions, for all groups of participants (two-tailed paired t-test, generative task participants: dimensionality, t-statistic 2.21, p-value 0.03; boundary, 6.21, 1e-7; volume, 9.57, 8e-13; robustness, 10.6, 2e-14. Maximum-likelihood task participants: dimensionality, 5.75, 5e-7; boundary, 4.79, 2e-6; volume, 10.8, 2e-14; robustness, 12.2, 2e-16). **b**: Data from individual ANNs (points), trained on the generative (orange) or maximum-likelihood (cyan) task. Network performance was always highest when evaluated against maximum-likelihood solutions, compared to generative solutions (all dots are above the identity line).

Put briefly, ANNs exhibited simplicity preferences only when trained to do so, whereas human participants exhibited them regardless of their instructions and training.

## Discussion

Simplicity has long been regarded as a key element of effective reasoning and rational decision-making. It has been proposed as a foundational principle in philosophy (Baker, 2022), psychology (Chater & Vitányi, 2003; Feldman, 2016), statistical inference (Balasubramanian, 1997; de Mulatier & Marsili, 2024; Grünwald, 2007; Gull, 1988; H. Jeffreys, 1939; MacKay, 1992; Wallace, 2005; Xie & Marsili, 2024), and more recently machine learning (Chaudhari et al., 2019; De Palma et al., 2019; Valle-Perez et al., 2019; Yang et al., 2022; but see Dubova et al., 2025 for a recent perspective on how highly complex models have recently challenged and complemented this traditional view across the sciences). Accordingly, various forms of simplicity preferences have been identified in human cognition (Gershman & Niv, 2013; Little & Shiffrin, 2009; Lombrozo, 2016; Pothos & Chater, 2002), such as a tendency to prefer smoother (simpler) curves as the inferred, latent source of noisy observed data (Genewein & Braun, 2014; Johnson et al., 2014), and visual perception related to grouping, contour detection, and shape identification (Feldman & Singh, 2006; Froyen et al., 2015; Wilder et al., 2016). However, despite the solid theoretical grounding of these studies, none attempted to define a quantitative notion of simplicity bias whose magnitude could be measured (as opposed to simply detected) in human behavior. In this work, we showed that simplicity preferences are closely related to a specific mathematical formulation of Occam’s razor, situated at the convergence of Bayesian model selection and information theory (Balasubramanian, 1997). This formulation enabled us to go beyond the mere detection of a preference for simple explanations for data and to measure precisely the strength of this preference in artificial and human participants under a variety of theoretically motivated conditions.

Our study makes several novel contributions. The first is theoretical: we derived a new term of the Fisher Information Approximation (FIA) in Bayesian model selection that accounts for the possibility that the best model is on the boundary of the model family (see details in Methods section A.2). This boundary term is important because it can account for the possibility that, because of the noise in the data, the best value of one parameter (or of a combination of parameters) takes on an extreme value. This condition is related to the phenomenon of “parameter evaporation” that is common in real-world models for data (Transtrum et al., 2015). Moreover, boundaries for parameters are particularly important for studies of perceptual decision-making, in which sensory stimuli are limited by the physical constraints of the experimental setup and thus reasoning about unbounded parameters would be problematic for observers. For example, imagine designing an experiment that requires participants to report the location of a visual stimulus. In this case, an unbounded set of possible locations (e.g., along a line that stretches infinitely far in the distance to the left and to the right) is clearly untenable. Our “boundary” term formalizes the impact of considering the set of possibilities as having boundaries, which tend to increase local complexity by reducing the number of local hypotheses close to the data (see Figure 1b).

The second contribution of this work relates to Artificial Neural Networks: we showed that these networks can learn to use or ignore simplicity preferences in an optimal way (i.e., according to the magnitudes prescribed by theory), depending on how they are trained. These results are different from, and complementary to, recent work that has focused on the idea that implementation of simple functions could be key to generalization in deep neural networks (Chaudhari et al., 2019; De Palma et al., 2019; Valle-Perez et al., 2019; Yang et al., 2022). Here we showed that effective learning can take into account the complexity of the hypothesis space, rather than that of the decision function, in producing normative simplicity preferences. On the one hand, these results do not seem surprising, because ANNs, and deep networks in particular, are powerful function approximators that perform well in practice on a vast range of inference tasks (Bengio et al., 2021). Accordingly, our ANNs trained with respect to the true generative solutions were able to make effective decisions, including simplicity preferences, about the generative source of a given set of observations. Likewise, our ANNs trained with respect to maximum-likelihood solutions were able to make effective decisions, without simplicity preferences, about the maximum-likelihood match for a given set of observations. On the other hand, these results also provide new insights into how ANNs might be analyzed to better understand the kinds of solutions they produce for particular problems. In particular, assessing the presence or absence of the kinds of simplicity preferences that we observed might help identify if and/or how well an ANN is likely to avoid overfitting to training data and provide more generalizable solutions.

The third, and most important, contribution of this work relates to human behavior: people tend to use particular forms of simplicity preferences when making decisions that implicate particular underlying computations. From a theoretical perspective, the difference between a Bayesian solution (i.e., one that includes the simplicity preferences) and a maximum-likelihood solution (i.e., one that does not include the simplicity preferences) to these tasks is that the latter considers only the single best-fitting model from each family, whereas the former integrates over all possible models in each family. This integration process is what gives rise to the simplicity bias, which, crucially, cannot emerge from simpler mechanisms such as sampling over different possible causes of the stimulus due to an unreliable sensory representation or a stochastic choice process (see Figure 2). Our finding that people, unlike ANNs, tend to use the Bayesian solution even when instructed to use the maximum-likelihood solution suggests that we naturally do not make decisions based on the single best or archetypical instance within a family of possibilities but rather integrate across that family. Put more concretely in terms of our task, when told to identify the shape closest to the data points, participants were likely uncertain about which exact location on each shape was closest. They therefore naturally integrated over the possibilities, which induces the simplicity preferences as prescribed by the Bayesian solution. These findings will help motivate and inform future studies to identify where and how the brain implements and stores these integrated solutions to relevant decision problems.

Another key feature of our findings that merits further study is the magnitude and variability of preferences exhibited by the human participants. On average, human sensitivity to each geometrical model feature was: 1) larger than zero, 2) at least slightly different from the optimal value (e.g., larger for dimensionality and robustness, smaller for volume), 3) different for distinct features and different participants, and 4) relatively insensitive to instructions and training. What is the source of this diversity? One hypothesis is that people may weigh more heavily the model features that are easier or cheaper to compute.

In our experiments, the most heavily weighted feature was model dimensionality. In our mathematical framework, this feature corresponds to the number of degrees of freedom of a possible explanation for the observed data and thus can be relatively easy to assess. By contrast, the least heavily weighted feature was model volume. This feature counts the number of distinguishable probability distributions contained within a mode and thus involves integrating over the whole model family. In other words, to count how many distinct states of the world can be explained by a certain hypothesis, one needs to enumerate them. This operation can be computationally costly (although for the models we used in our tasks, this number is related directly to the length of the lines drawn on the screen measured in units of the standard deviation of the noise and thus could be fairly straightforward for our participants to infer, see Methods section A.4).

The other two terms, boundary and robustness, are intermediate in terms of human weighting and computational difficulty. They are harder to compute than dimensionality, because they depend on the data and on the properties of the model at the maximum-likelihood location. However, they are also simpler than the volume term, because they are local quantities that do not require integration over the whole model manifold. This intuition leads to new questions about the relationship between the complexity of explanations being compared and the complexity of the decision-making process itself, calling into question notions of bounded rationality and diminishing returns in optimal inference (Tavoni et al., 2019, 2022). Answering such questions is beyond the scope of the present work but merits further study.

A different, intriguing future direction is a comparison with other formal approaches to the emergence of simplicity that can lead to different predictions. Recent studies have argued that Jeffrey’s prior (upon which our geometric approach is based) could give an incomplete picture of the complexity of a class of models that occur commonly in the natural sciences, which contain many combinations of parameters that do not affect model behavior, and proposed instead the use of data-dependent priors (Mattingly et al., 2018; Quinn et al., 2022). The two methods lead to different results, especially in the data-limited regime (Abbott & Machta, 2023). It would be useful to understand the relevance of these differences to human and machine decision-making.

Our task design and analyses had several limitations that can be addressed in future studies. We used only four task variants, which limits the generalizability of our findings (although note that because the boundary and robustness penalties depend on the model and on the data, they both assumed a continuum of values through the tasks, and the boundary penalty was relevant for all task types). We also considered only a fixed sample size *N*, conditions for which the FIA for the models involved could be computed analytically, and independent trials that ignore sequential aspects of repeated decisions. Extending our approach to relax these assumptions, or to include conditions in which the volume or dimensionality of the models is allowed to vary more freely, would provide further insights into when and how people exhibit simplicity preferences when making decisions.

In summary, our work demonstrates the direct, quantitative relevance of formal notions of model complexity to human behavior. By relying on a combination of theoretical advances, computational modeling, and behavioral experiments, we have established a novel set of normative reference points for decision-making under uncertainty. These findings open up a new arena in which human cognition could be measured against optimal inferential processes, potentially leading to new insights into the constraints affecting information processing in the brain.

## Data availability

All experimental data collected in this work is available at doi:10.17605/OSF.IO/R6D8N.

## Code availability

All data and code needed to reproduce the experiments (including running the online psychophysics tasks and training and testing the neural networks), and to analyze the data and produce all figures is available at doi:10.17605/OSF.IO/R6D8N.

## Ethics

Human participant protocols were approved and determined to be Exempt by the University of Pennsylvania Internal Review Board (IRB protocol 844474). Participants provided consent on-line before they began the task.

## Acknowledgements

We thank Kamesh Krishnamurthy for discussions and acknowledge the financial support of R01 NS113241 (EP), R01 EB026945 (JIG and VB), IIS-2145164 (PC), CCF-2212519 (PC) as well as a hardware grant from the NVIDIA Corporation (EP). The HPC Collaboration Agreement between SISSA and CINECA granted access to the Marconi100 and Leonardo clusters. VB was supported in part by the Eastman Professorship at Balliol College, University of Oxford.

## Author contribution

Conceptualization: EP VB JG. Methodology: EP SL PC VB JG. Software: EP SL. Formal analysis: EP SL. Investigation: EP SL. Resources: EP VB JG. Data curation: EP SL. Writing - original draft: EP JG. Writing - editing and reviewing: EP SL PC VB JG. Supervision: VB JG. Project administration: JG. Funding acquisition: VB JG.

## Appendix A

### Methods

#### A.1 Noise-Integration-Noise (NIN) ideal observer

To formalize the intuition around Occam’s razor emerging from a process of integration over latent causes, we define an ideal observer that can perform the model-selection tasks described in the main text based on a simple set of heuristics. The observer has three degrees of freedom: 1) the intensity of sensory noise, 2) the extent to which it performs integration over latent causes, and 3) the intensity of the choice noise. We call this the Noise-Integration-Noise (NIN) observer.

In particular, we assume that some empirical data 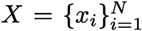 are generated from one of the models described in section A.4. Briefly, each datapoint *x*_*i*_ ∈ ℝ^2^ is sampled independently with

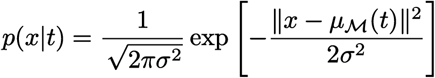

where *σ* is fixed and known (in practice, *σ* = 1 for our simulations), ∥ · ∥ is the Euclidean norm in ℝ^2^, and the (smooth) function *µ*_M_ : [0, 1] → ℝ^2^ is specific to the model ℳ generating the data. For instance, in the example shown in Figure 1 in the main text (the Dimensionality task), *µ*_ℳ_ (*t*) = (0, 0) for the model on the bottom (the dot) and *µ*_ℳ_ (*t*) = ((*t* − 1)*/*2, 1) for the model on top (the line).

For data generated according to the procedure above, the ideal observer starts by computing the average of the empirical data

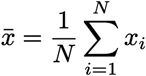

This representation of the data is corrupted by normally distributed sensory noise of intensity *ρ*:

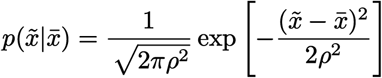

Based on this (noisy) internal representation of the data, the participant computes an approximate Bayesian posterior, based on computing the evidence in a restricted neighborhood of the maximum-likelihood point:

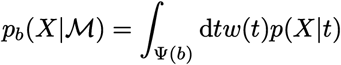

where *w* is uniform over Ψ, and the neighborhood Ψ(*b*) is defined as a function of the integration parameter *b* in such a way that the observer will:

1. perform full Bayesian integration for *b* = 1,
2. take into account only the maximum-likelihood point for *b* = 0, so that 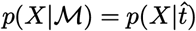, and
3. will perform an operation that interpolates between these two extremes for intermediate values of *b*. This behavior is obtained by setting

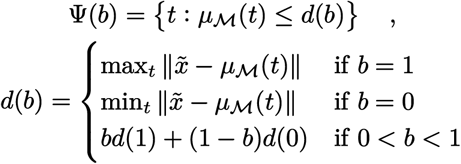

In practice, the observer considers only the portion of the model contained within a circle of radius *d*(*b*) centered on the data, with the radius set such that: 1) for *b* = 0, the circle will touch the model only at the point that is closest to the data; 2) for *b* = 1, the circle will include all of the model; and 3) for intermediate values, the circle will interpolate linearly between these two cases.

Finally, the participant picks a model sampling from the following distribution:

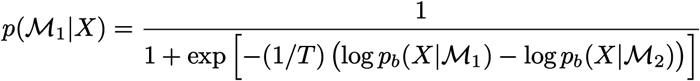

Where the “temperature” parameter *T* controls the level of choice noise. This procedure corresponds to sampling from the approximate Bayesian posterior when *T* = 1, picking the model with the highest posterior for *T* → 0, and picking a model at random for *T* → ∞.

##### A.1.1 Model-free analysis of simplicity bias for the NIN observer

We performed an elementary, model-free assessment of the simplicity bias exhibited by the NIN observer as a function of the observer’s parameters. For a given configuration of the observer, we simulated 10000 trials for each of the task types described in the main text. We then quantified the choice bias by fitting a sigmoid psychometric curve (as a function of *y*, the vertical coordinate) to data falling in a restricted region in the center of the data space (−0.1 *< x <* 0.1, 0 *< y <* 1):

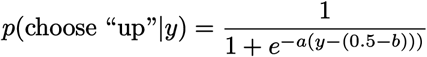

where *a* governs the shape (slope) of the psychometric curve, and *b* governs the bias that here is defined as the distance between the decision boundary and the midpoint between the models (*x* = 0, *y* = 0.5). The sign of the bias is taken such that a positive bias pushes the decision boundary towards the more complex model (this is the direction expected from Occam’s razor), and a negative bias has the opposite effect.

#### A.2 Derivation of the boundary term in the Fisher Information Approximation

Here we generalize the derivation of the Fisher Information Approximation (FIA) given by Balasubramanian (1997) to the case where the maximum-likelihood solution for a model lies on the boundary of the parameter space. This generalization is important because it relies on more realistic assumptions than existing approaches. In particular, existing approaches typically assume that the maximum-likelihood solution is in the interior of the parameter space of a given model. In contrast, models are just approximations of the true processes in the real world that generated a given set of observations, implying that those observations may fall outside of the range that can be expected based on the parameter space of a given model. In these cases (or even when the observations are based on samples generated by the model but are corrupted by noise to fall outside of the range implied by most values of the model’s parameters space), the maximum-likelihood solution for that model, given those data, may fall on the boundary of the model’s parameter space. To account for this condition, we extended the FIA to deal with the simple case of a linear boundary in parameter space. When the maximum-likelihood solution is on such a boundary, an additional penalty term appears in the FIA, which we denote “boundary” (see Figure 1d in the main text).

Apart from the more general assumptions, the following derivation follows closely the original one, with some minor notational changes. This derivation appeared in preliminary form in Piasini et al. (2021a).

##### Sketch of the derivation

Our derivation below uses a well-known approximation method to yield a closed-form expression for the model evidence in the limit of large sample size (large *N*). The evidence is the integral of the likelihood over all possible parameter values, weighted by the prior. Up to constant terms, this integral can be written in the form ∫d*ϑ* exp [−*Nψ*(*ϑ, X*)], where *ψ*(*ϑ, X*) is a function that depends on the data *X* and on a location in parameter space *ϑ*. In a nutshell, the approximation consists in noticing that, as *N* increases, the integral will come to be dominated by the behavior of the integrand within a small neighborhood of the maximum-likelihood point 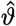 in parameter space (which is itself dependent on the data: 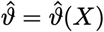. In this neighborhood, we rewrite the integrand by Taylor-expanding *ψ*(*ϑ, X*) around the maximum-likelihood point. By inspecting the resulting series, we drop the terms of the expansion that decrease in magnitude when *N* grows, keeping only those that dominate in that regime. The end result is a Gaussian integral, which we can solve in closed form to yield the Fisher Information Approximation. This procedure is the same whether the maximum-likelihood point is in the interior of or on the boundary of the parameter space; the difference is that with a few technical passages we can show that, in the latter case, an additional term appears in the solution to the Gaussian integral. Intuitively, this extra term is due to the fact that when the maximum-likelihood point 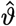 is on the boundary of the parameter space the gradient of the likelihood in 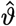 is not zero.

##### A.2.1 Set-up and hypotheses

The problem we consider here is that of selecting between two models (say ℳ_1_ and ℳ_2_), after observing empirical data 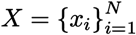. *N* is the sample size and ℳ_1_ is assumed to have *d* parameters, collectively indexed as *ϑ* taking values in a compact domain Θ. As a prior over *ϑ* we take Jeffreys prior:

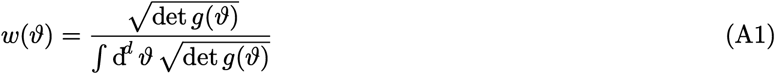

where *g* is the (expected) Fisher Information of the model ℳ_1_:

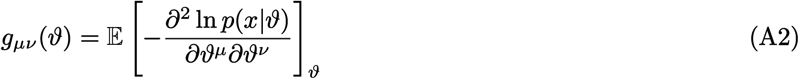

The Bayesian posterior

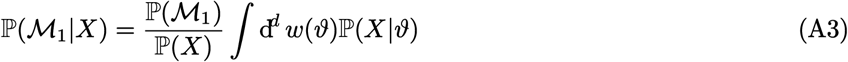

then becomes, after assuming a flat prior over models and dropping irrelevant terms:

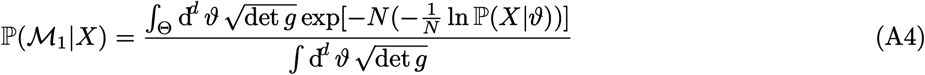

Just as in Balasubramanian (1997), we now make a number of regularity assumptions: 1) ln ℙ (*X*|*ϑ*) is smooth; 2) there is a unique global minimum 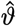 for ln ℙ (*X*|*ϑ*); 3) *g*_*µν*_(*ϑ*) is smooth; 4) 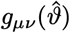 is positive definite; 5) Θ ⊂ ℝ^*d*^ is compact; and 6) the values of the local minima of ln ℙ (*X*|*ϑ*) are bounded away from the global minimum by some *ϵ >* 0. Importantly, unlike in Balasubramanian (1997), we do not assume that 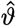 is in the interior of Θ.

###### The shape of Θ

Because we are interested in understanding what happens at a boundary of the parameter space, we add a further assumption that, while being not very restrictive in spirit, allows us to derive a particularly interpretable result. In particular, we assume that Θ is specified by a single linear constraint of the form:

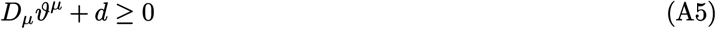

Without loss of generality, we also take the constraint to be expressed in Hessian normal form, namely, ∥*D*_*µ*_∥ = 1. For clarity, note this assumption on the shape of Θ is used only from subsubsection A.2.3 onward.

##### A.2.2 Preliminaries

We now proceed to set up a low-temperature expansion of Equation A4 around the saddle point 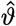. We start by rewriting the numerator in Equation A4 as:

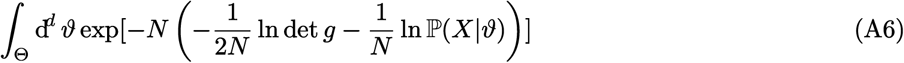

The idea of the FIA is to expand the integrand in Equation A6 in powers of *N* around the maximum likelihood point 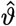. To this end, we define three useful objects:

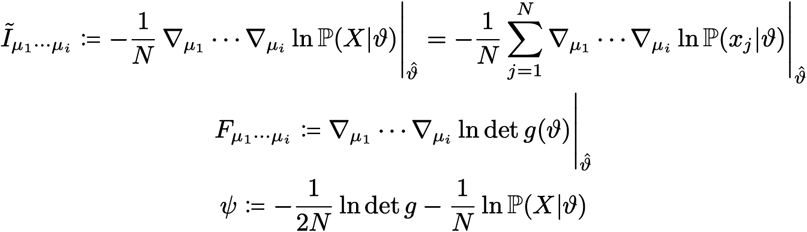

We immediately note that:

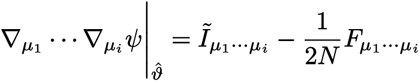

which is useful to compute

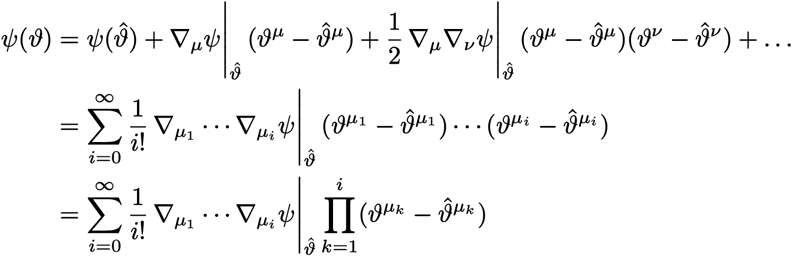

It is also useful to center the integration variables by introducing

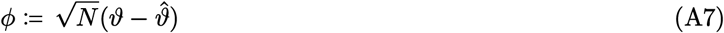

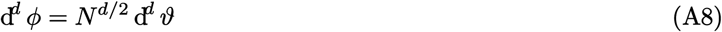

so that

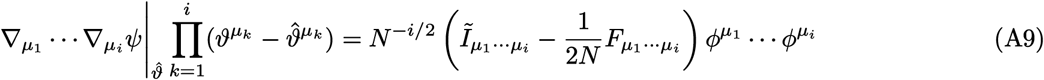

and Equation A6 becomes:

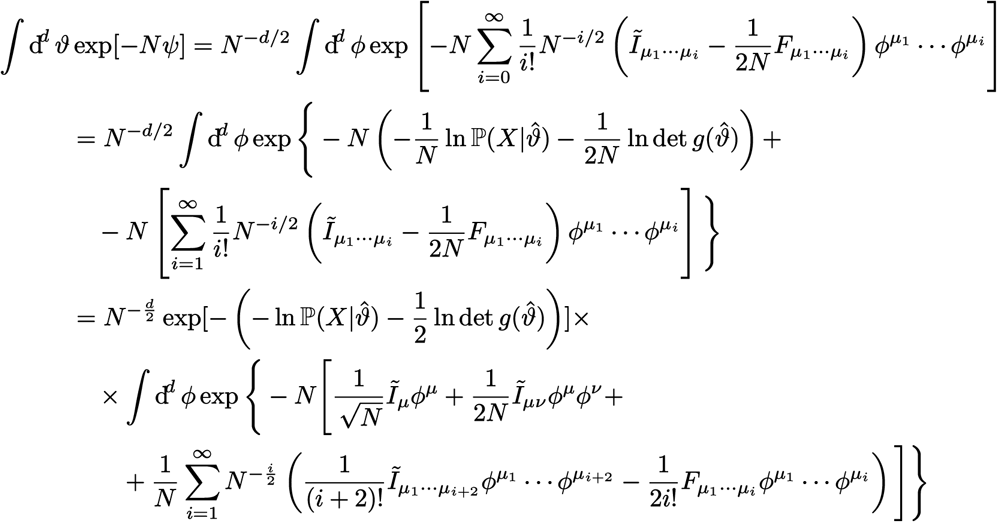

Therefore,

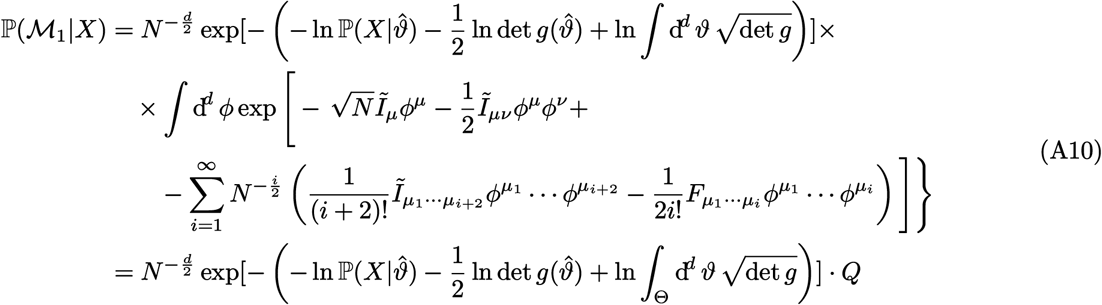

where

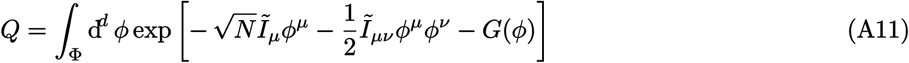

and

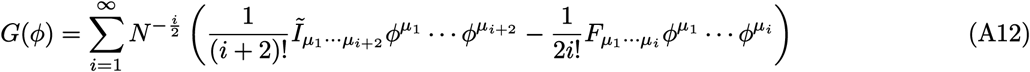

where *G*(*φ*) collects the terms that are suppressed by powers of *N* .

Our problem has been now reduced to computing *Q* by performing the integral in Equation A11. Now our assumptions come into play for the key approximation step. For the sake of simplicity, assuming that *N* is large we drop *G*(*φ*) from the expression above, so that *Q* becomes a simple Gaussian integral with a linear term:

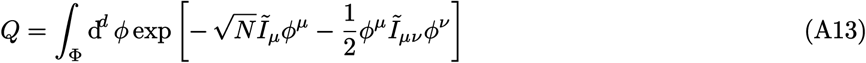

##### A.2.3 Choosing a good system of coordinates

Consider now the Observed Fisher Information at the maximum likelihood, *Ĩ*_*µν*_ . As long as it is not singular, we can define its inverse Δ_*µν*_ = (*Ĩ*_*µν*_)^−1^. If *Ĩ*_*µν*_ is positive definite, then the matrix representation of *Ĩ*_*µν*_ has a set of *d* positive eigenvalues, which we denote by 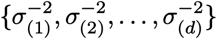. The matrix representation of Δ^*µν*^ has eigenvalues 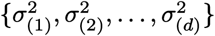, and is diagonal in the same choice of coordinates as *Ĩ*_*µν*_ . We denote by *U* the (orthogonal) diagonalizing matrix; i.e., *U* is such that

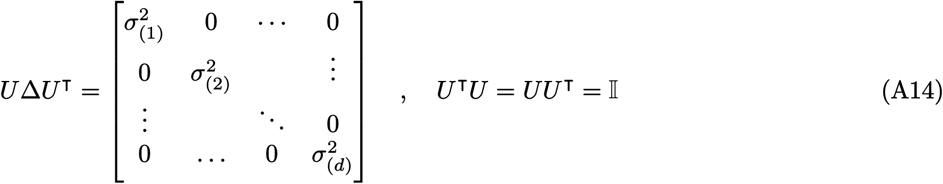

We define also the matrix *K* as the product of the diagonal matrix with elements 1*/σ*_(*k*)_ along the diagonal and *U* :

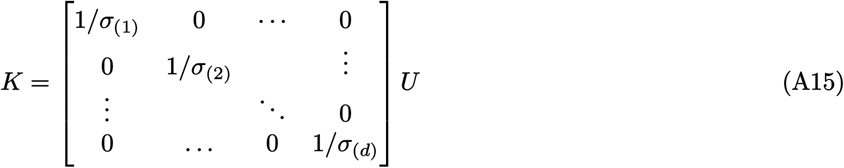

Note that

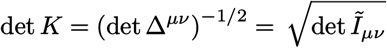

and that *K* corresponds to a sphering transformation, in the sense that

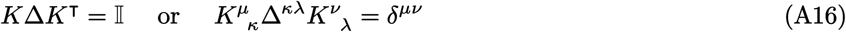

and therefore, if we define the inverse we have

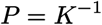

We have

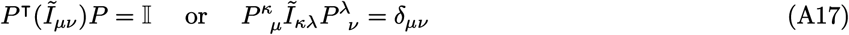

We can now define a new set of coordinates by centering and sphering, as follows:

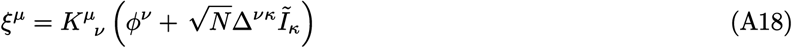

Then,

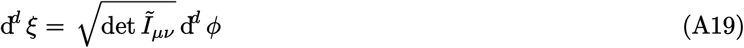

and

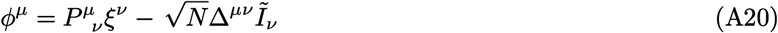

In this new set of coordinates,

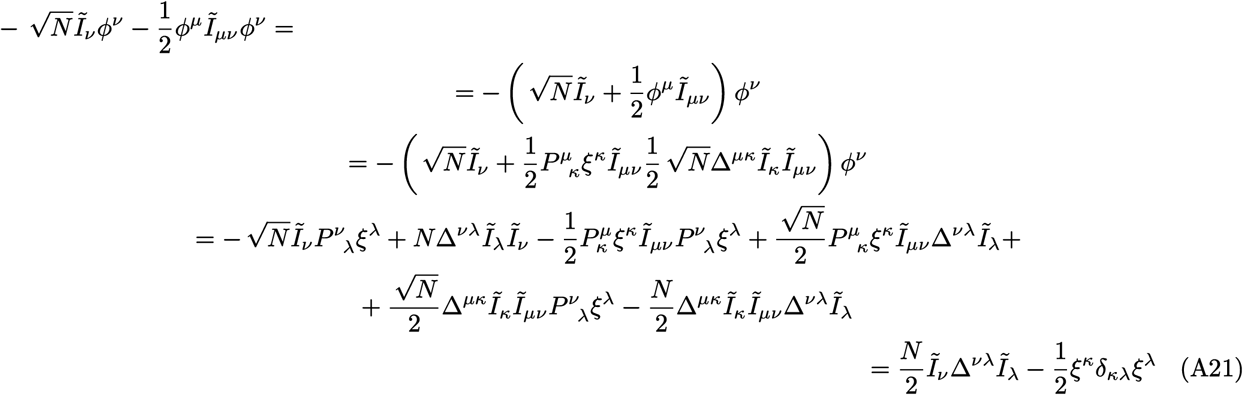

where we have used Equation A17 as well as the fact that Δ^*µν*^ = Δ^*νµ*^ and that Δ^*µκ*^*Ĩκν* = *δ*^*µ*^_*ν*_ by definition.

Therefore, putting Equation A19 and Equation A21 together, Equation A13 becomes

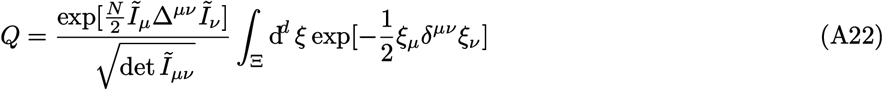

The problem is reduced to a (truncated) spherical gaussian integral, where the domain of integration Ξ will depend on the original domain Θ but also on *Ĩ*_*µ*_, *Ĩ*_*µν*_ And 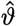. To complete the calculation, we now need to make this dependence explicit.

##### A.2.4 Determining the domain of integration

We start by combining Equation A7 and Equation A20 to yield:

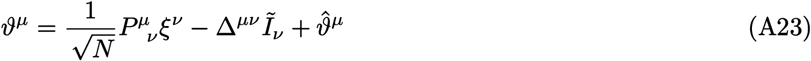

By substituting Equation A23 into Equation A5 we get

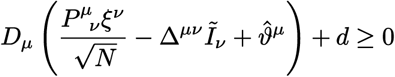

which we can rewrite as

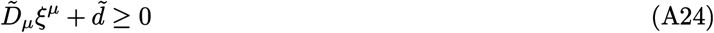

with

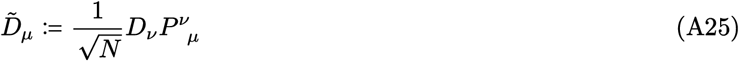

and

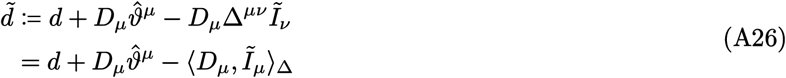

where by ⟨·, ·⟩_Δ_ we mean the inner product in the inverse observed Fisher information metric. Note that whenever *Ĩ*_*µ*_ is not zero, it will be parallel to *D*_*µ*_. Indeed, by construction of the maximum-likelihood point 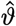, the gradient of the log likelihood can be orthogonal to the boundary only at 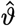, where it points towards the outside of the domain. Therefore *Ĩ*_*µ*_, which is defined as minus the gradient, will point inward. At the same time, *D*_*µ*_ will also always point toward the interior of the domain because of the form of the constraint we have chosen in Equation A5. Because by assumption ∥*D*_*µ*_∥ = 1, we have that

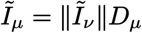

and

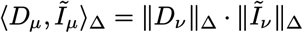

so that

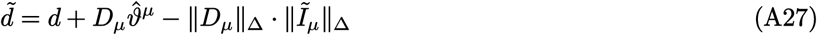

Now, the signed distance of the boundary to the origin in *ξ*-space is

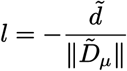

where the sign is taken such that *l* is negative when the origin is included in the integration domain. But noting that

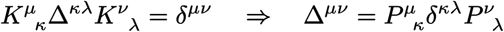

we have

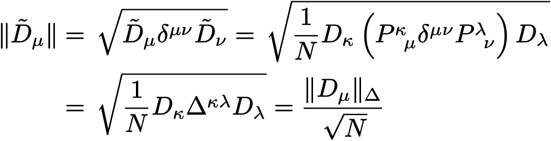

and therefore

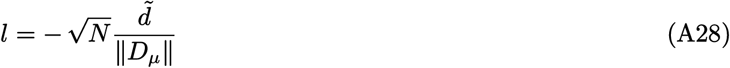

Finally, by plugging Equation A27 into Equation A28 we obtain

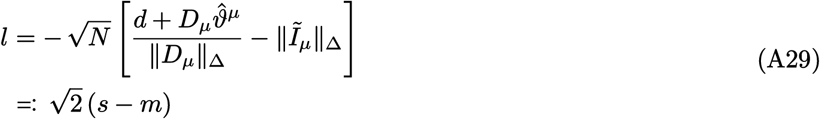

where *m* and *s* are defined for convenience like so:

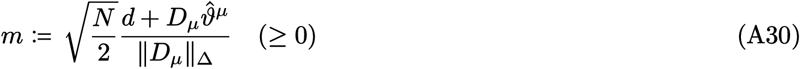

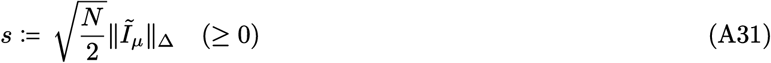

We note that *m* is a rescaled version of the margin defined by the constraint on the parameters (and therefore is never negative by assumption), and *s* is a rescaled version of the norm of the gradient of the log likelihood in the inverse observed Fisher metric (and therefore is nonnegative by construction).

##### A.2.5 Computing the penalty

We can now perform a final change of variables in the integral in Equation A22. We rotate our coordinates to align them to the boundary, so that

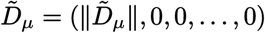

Note that we can always do this as our integrand is invariant under rotation. In this coordinate system, Equation A22 factorizes:

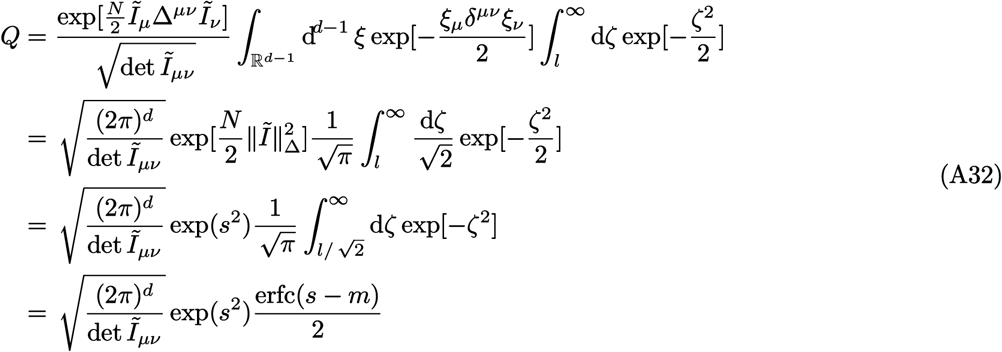

where erfc(·) is the complementary error function (Abramowitz & Stegun, 1972, section 7.1.2).

Finally, plugging Equation A32 into Equation A10 and taking the log, we obtain the extended FIA:

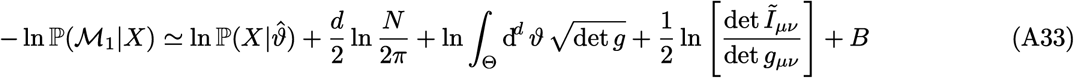

where

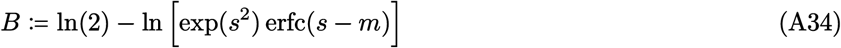

can be interpreted as a penalty arising from the presence of the boundary in parameter space.

##### A.2.6 Interpreting the penalty

We now take a closer look at Equation A34. One key observation is that, by construction, at most one of *m* and *s* is ever nonzero. In the interior of the manifold, *m >* 0 by definition, but *s* = 0 because the gradient of the likelihood is zero at 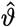. On the boundary, *m* = 0 by definition, and *s* can be either zero or positive.

###### Interior of the manifold

When 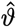 is in the interior of the parameter space Θ, then *Ĩ*_*µ*_ = 0 ⇒ *s* = 0, and Equation A34 simplifies to

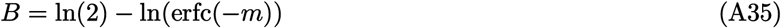

but because *N* is large we have *m* ≫ 0, erfc(− *m*) → 2 and *B* → 0, so our result passes the first sanity check: we recover the expression in Balasubramanian (1997).

###### Boundary of the manifold

When 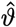 is on the boundary of Θ, *m* = 0 and *s* ≥ 0. Equation A34 becomes

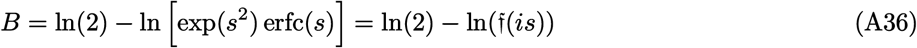

where f is the Faddeeva function (Abramowitz & Stegun, 1972, p. 7.1.3):

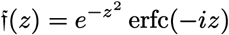

This function is tabulated and can be computed efficiently. However, it is interesting to analyze its limiting behavior, as follows.

As a consistency check, when *s* is small we have at fixed *N*, to first order:

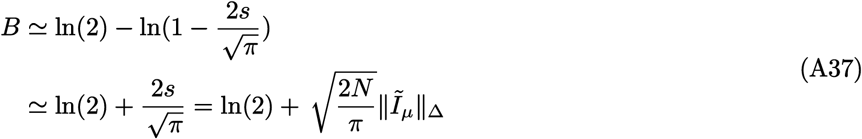

and *B* = ln(2) when *Ĩ*_*µ*_ = 0, as expected.

However, the real case of interest is the behavior of the penalty when *N* is assumed to be large, which is consistent with the fact that we derived Equation A32 as an asymptotic expansion of Equation A11. In this case, using the asymptotic expansion for the Feddeeva function (Abramowitz & Stegun, 1972, section 7.1.23):

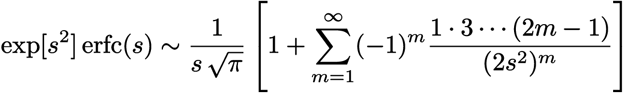

To leading order, we obtain

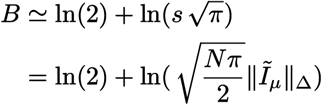

which we can rewrite as

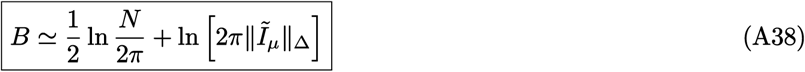

We can summarize the above by saying that a new penalty term of order ln *N* arose due to the presence of the boundary. Interestingly, comparing Equation A38 with Equation A33 we see that the first term in Equation A38 is analogous to counting an extra parameter dimension in the original Fisher Information Approximation.

#### A.3 Human psychophysics

The behavioral task required participants to view a screen showing two curves (one on the upper half, the other on the lower half of the screen) and 10 dots and decide, based on different instructions (see below for details), which curve was the more likely source of the observed dots. There were four task types that differed in terms of the shapes of the curves, corresponding to the different terms of the FIA (see Figure 1 in the main text and Figure B.1): *dimensionality, boundary, volume*, and *robustness*. In each case, the curves represent two parametric statistical models of the form:

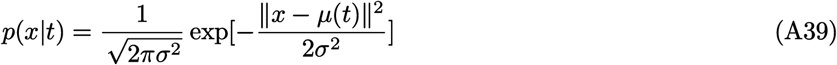

where *x* is a location on the 2D plane visualized on the screen, and *µ*(*t*), *t* ∈ [0, 1] is a parametrization of the curve. In other words, the curves represent Gaussians of unit isotropic variance whose mean *µ* can be located at any point along them. The dots shown to the participants were sampled iid from one of the two models, selected at random with uniform probability. The location of the true mean of the Gaussian generating the dots (i.e., the value of *t* in the expression above) was sampled randomly from Jeffreys’ prior for the selected model. All dots shown within a trial come from the same distribution (same model and same true mean). In the “generative” version of the task, the participants had to report which curve (model) the dots are more likely to come from. In the “maximum-likelihood” version, the participants had to report which curve was closest to the empirical centroid of the dot cloud. In both versions of the task, they pressed the “up” or “down” keys on their keyboard to select the curve in the upper or lower part of the screen, respectively.

Each model pairing was designed to emphasize a different term of the FIA. In the dimensionality variant, models have different dimensionality (*d* = 0 for the point and *d* = 1 for the line). In the boundary variant, both models have the same dimensionality and volume and are both flat so that their robustness terms are always identically zero. However, they are oriented such that, for ambiguous data falling around the midpoint between the two models, the influence of the boundary of the vertical model is stronger than that of the horizontal model. In the volume variant, models have the same dimensionality but different volumes (length). In the robustness variant, models have the same dimensionality and volume, but their curvature is such that one of them bends away from the region of data space that is more likely to contain ambiguous stimuli, whereas the other bends around it (and therefore the robustness term for these models has opposite sign for data points that fall in that region).

A single run of the task consisted of a brief tutorial followed by 500 trials, divided in 5 blocks of 100 trials each. For each trial, the chosen curve pairing was presented, randomly flipped vertically to dissociate a fixed preference for one of the two models from a fixed preference for reporting “up” or “down”. At the end of each block, the participant received feedback on their overall performance during that block. Participants received a fixed compensation for taking part in the experiment.

We report how we determined our sample size, all data exclusions (if any), all manipulations, and all measures in the study. We ran both experiments (generative and maximum-likelihood) on the online platform Pavlovia (pavlovia.org). For each task type, we collected data from at least 50 participants who passed a pre-established performance threshold (60% correct for the robustness task variant and 70% correct for the other variants; the sample size and the thresholds were chosen based on pilot data and were fixed at preregistration (Piasini et al., 2020, 2021b, 2022)). We discarded the data collected from all other participants. For the generative task, the final dataset included 52 participants for the robustness task variant and 50 participants for each of the other task variants. For the maximum-likelihood task, the final dataset included 51 participants for the dimensionality task variant and 50 participants for each of the others.

#### A.4 Detailed model definitions and computation of FIA terms

In this section, we report the detailed mathematical form of the models we used for the psychophysics experiment. Each model is defined by specifying the form of the function *µ* in Equation A39. Given this function, we then derive the analytical solution to the maximum-likelihood problem for any value of 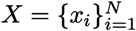, and finally the expressions for the likelihood (*L*), dimensionality (*D*), boundary (*B*), volume (*V*), and robustness (*R*) terms in the FIA for the model pairings we use in the experiment.

We also show that the (expected) Fisher information is constant for all models considered:

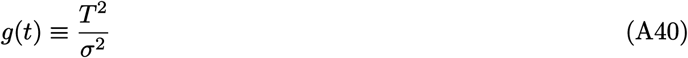

so that Jeffreys prior is simply the uniform probability distribution over the [0, 1] interval:

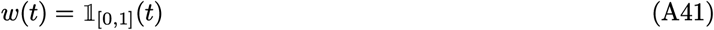

##### A.4.1 Fisher information and robustness term for curved exponential families

In the following, we compute the observed Fisher information for each of our models. To do so, it is convenient to have a general expression for the Hessian of the log likelihood and for the observed and expected Fisher information for curved exponential families.

The general form of a curved exponential family is:

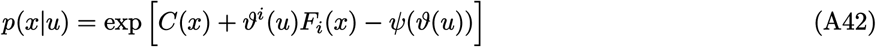

where *ϑ*(*u*) : ℝ^*d*^ → ℝ^*k*^, *k* ≥ *d*, is a smooth parametrization. The Hessian of the log-likelihood is:

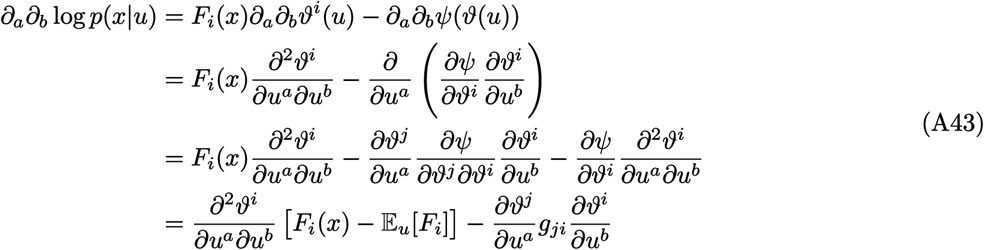

where we note that *g*_*ij*_ = −Cov_*u*_[*F*]_*ji*_ (remember that by *g*_*ij*_ we indicate the Fisher information of the ambient family). Therefore, the (expected) Fisher information is:

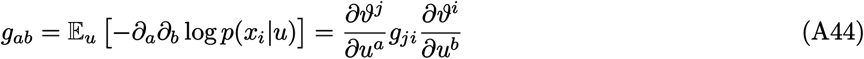

and the oberved Fisher information is:

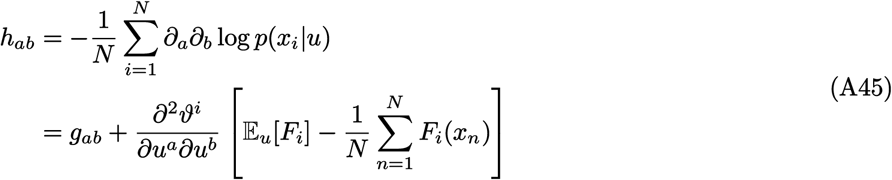

As a corollary, we note that *h*_*ab*_ = *g*_*ab*_ whenever *ϑ*(·) is an affine transformation, that is when

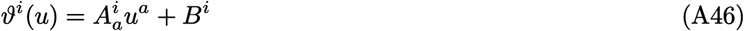

For some constant 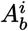 and 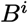. In this case (which corresponds to autoparallel submanifolds in the exponential connection, (Amari & Nagaoka, 2000, Theorem 1.1)), the robustness term in the FIA is identically zero:

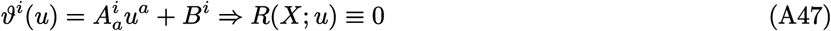

##### A.4.2 General properties of curved 2D Gaussian models

Our models of interest, defined through Equation A39, are a special case of curved exponential families. They are all submanifolds of the same, larger model — the 2-dimensional exponential family of 2D Gaussian distributions with known isotropic covariance and unknown center. We call this larger family the *ambient family S* ⊃ ℳ, composed by all probability distributions whose density is of the form:

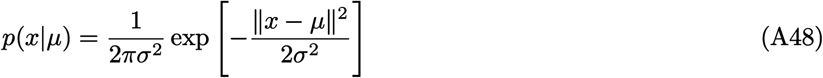

We can reduce Equation A39 to the notation of Equation A42 by noting that

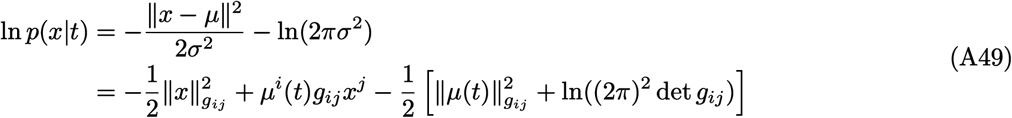

where we indicate by *g*_*ij*_ the Fisher information of the ambient family *S*:

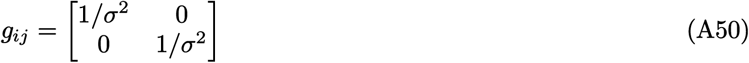

By comparing Equation A49 with Equation A42, we see that

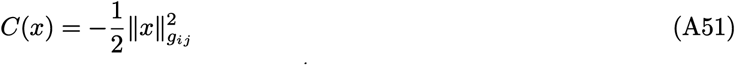

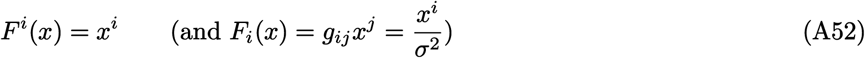

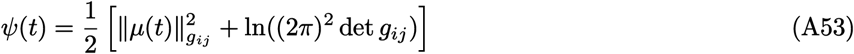

and that *µ*(*t*) plays the role that *ϑ*(*u*) played in Equation A42.

We can now compute the expected and observed Fisher information for our models by specializing Equation A44 and Equation A45:

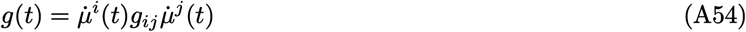

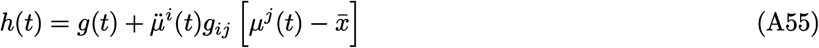

Where 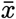 is the empirical centroid of the dataset *X*,

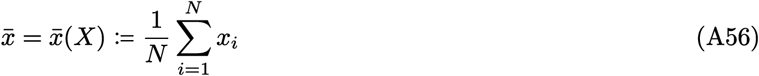

and *g* and *h* have no indices, because they are scalar functions of *t*.

We note then that *g*(*t*) is simply the squared Euclidean norm of the vector 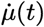 divided by *σ*^2^. In other words, the geometry of ℳ coincides, up to scaling by *σ*^2^, with the Euclidean geometry of the plane curve *µ*(*t*). This very convenient fact is a consequence of the particularly simple noise model we have assumed (Gaussian with known isotropic covariance).

###### Model volume

The volume of a model described by *µ*(·) is

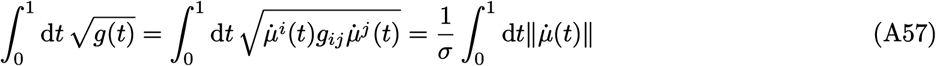

In other words, it is simply the length of the curve *µ*(·) measured in units of *σ*.

###### Likelihood gradient and maximum-likelihood point

In the following, we will indicate the log-likelihood function for a model by

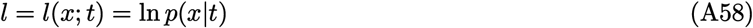

In order to find the maximum-likelihood point for our models, it is convenient to write a general expression for the score function (the derivative of the log likelihood with respect to the parameter). We start by noting that

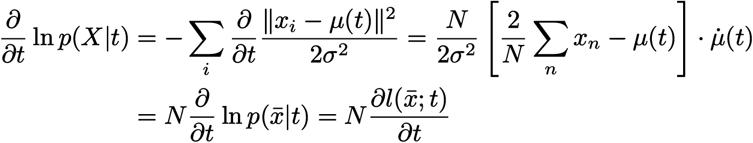

Therefore, to find the maximum-likelihood point 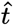 for a certain *X* we can simply solve the corresponding one-sample (*N* = 1) case for the centroid 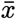. We can also write the rescaled likelihood gradient (which appears in the FIA as *I*_*µ*_) as

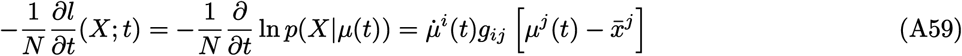

If we interpret 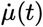 as the tangent vector to *µ* in *t*, we see that away from model boundaries this equation expresses the familiar condition that the maximum-likelihood point (where ∂*l/*∂*t* = 0) is the (Euclidean) orthogonal projection of 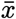 onto the model manifold. Again, this convenient property is a consequence of assuming isotropic Gaussian noise.

##### A.4.3 Horizontal model

This model, used in the Dimensionality, Boundary, and Volume tasks, is defined as

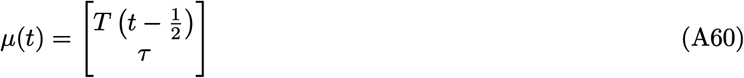

It is immediately evident that this model has volume (length) *T/σ*. The “base” model corresponds to *T* = 1, *τ* = 0, and the model type called “horizontal” is defined with *T* = 3, *τ* = 1.

Because

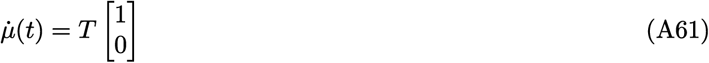

and following Equation A54 and Equation A55, the observed and expected Fisher information coincide and are given by

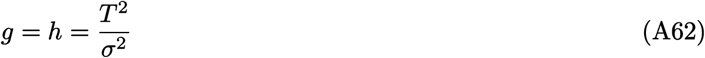

Given a centroid *X* = [*X*^1^, *X*^2^]^⊺^, the rescaled likelihood gradient is (from Equation A59)

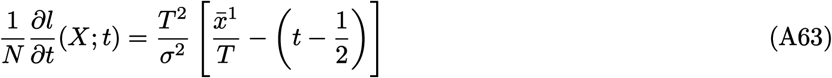

and the maximum-likelihood point 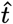 is

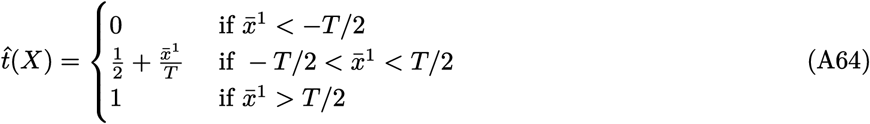

All the FIA terms can be computed in closed form from these expressions:

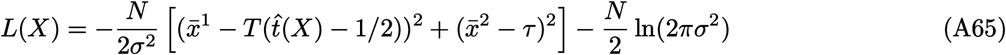

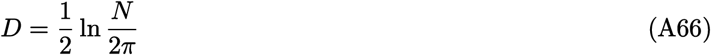

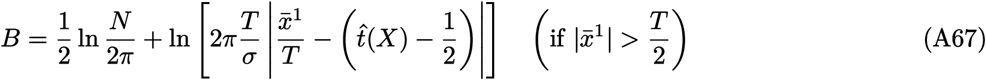

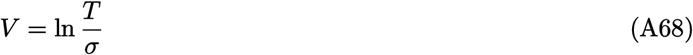

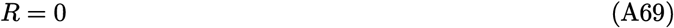

##### A.4.4 Vertical model

This model, used the Boundary task, is just a rotated and translated version of the horizontal model. It is defined as

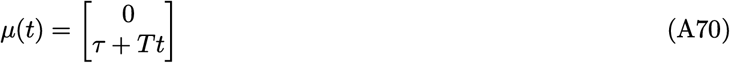

where we keep *T* and *τ* as arbitrary parameters for notational clarity, although in practice they are both fixed to 1 in our study. From the definition, it follows that

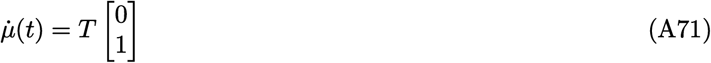

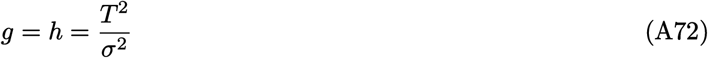

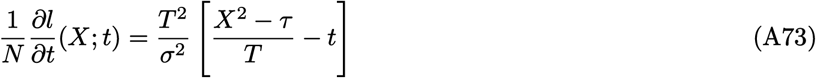

and

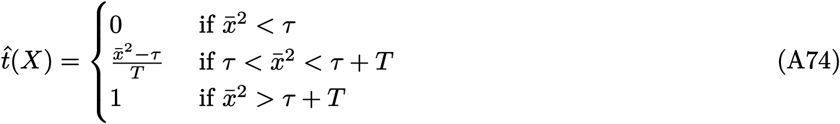

so that the FIA terms can be written as

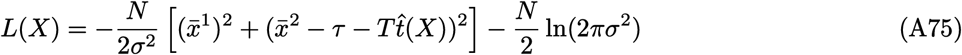

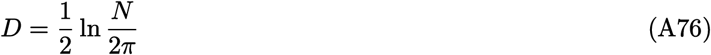

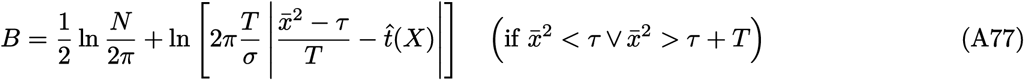

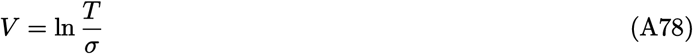

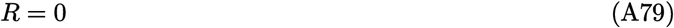

##### A.4.5 Circular-arc model

This model, used in the Robustness task, is constituted by an arc of a circle, and is defined as

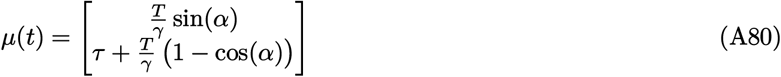

where

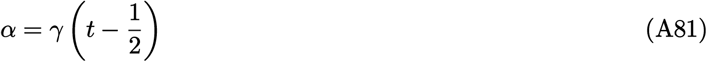

and *γ* is a positive constant. Concretely, in the experiments we fixed *γ* = (3*/*5)*π* and *T* to the value determined below for the rounded model type (Equation A99). The radius of the circle is *r* = *T/γ*, and the y-coordinate of the center is *τ* + *r*. The tangent vector 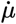 and the acceleration vector 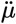 in *t* are, respectively:

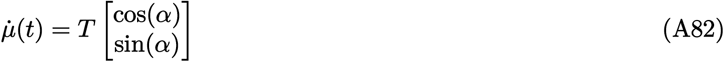

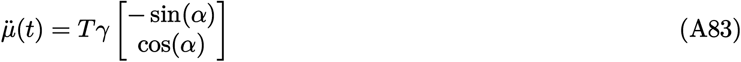

so that, by substitution in Equation A54,

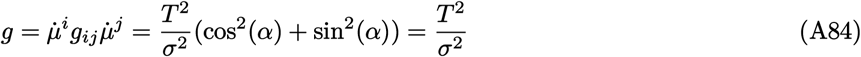

and by substitution in Equation A55

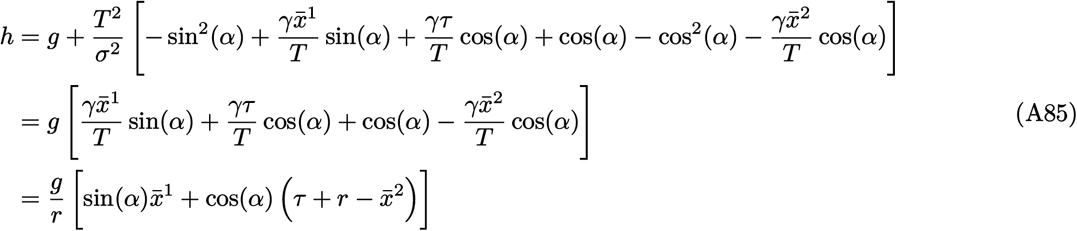

The rescaled likelihood gradient is (from Equation A59)

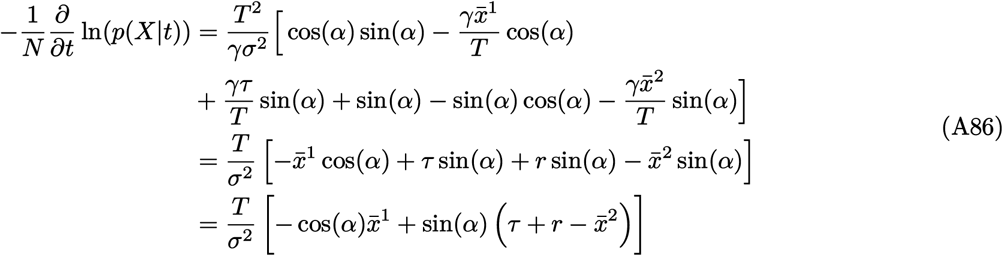

(note that *h* can also be obtained by differentiating this last expression).

To compute the FIA, we need the maximum-likelihood projection. As for the other models, this projection is defined piecewise because of the presence of model boundaries. To properly partition the plane, we need to define first the equation for the line intersecting the model perpendicularly at *t*:

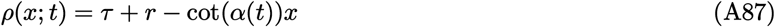

With this definition, the maximum-likelihood point is

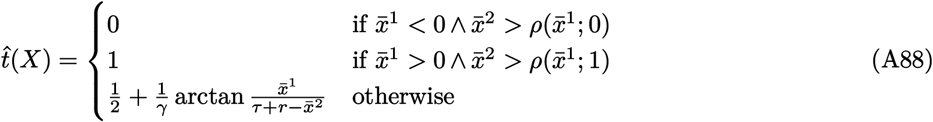

and therefore the FIA terms are:

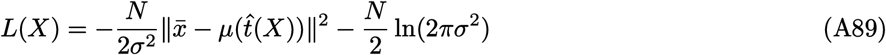

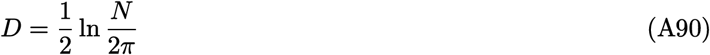

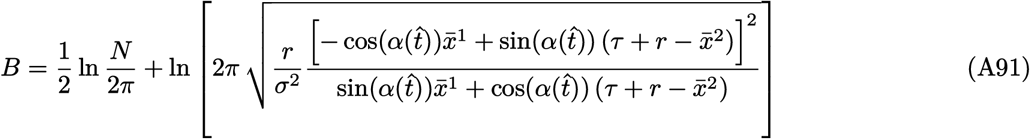

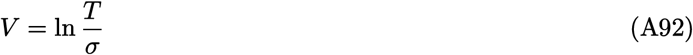

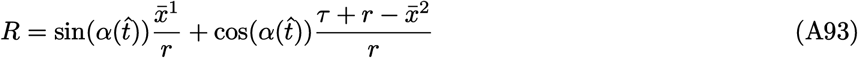

where the value given for *B* is relevant only when 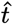 is either 0 or 1. Note that, because of the shape of the model and the presence of the boundary, there are regions of the data space such that the log-likelihood function at the maximum-likelihood point will not be concave. These regions represent a complete breakdown of the FIA, but they are not a problem in practice because the approximation holds in the region of data space that is relevant for the experiments (see Figure B.1).

##### A.4.6 Rounded model

This model, also used in the Robustness task, is a circular arc (like the “circular” model described above) with two straight arms attached on either side. The ratio of the length of the circular section of the model over its total length is defined as a parameter *f* . The model definition is

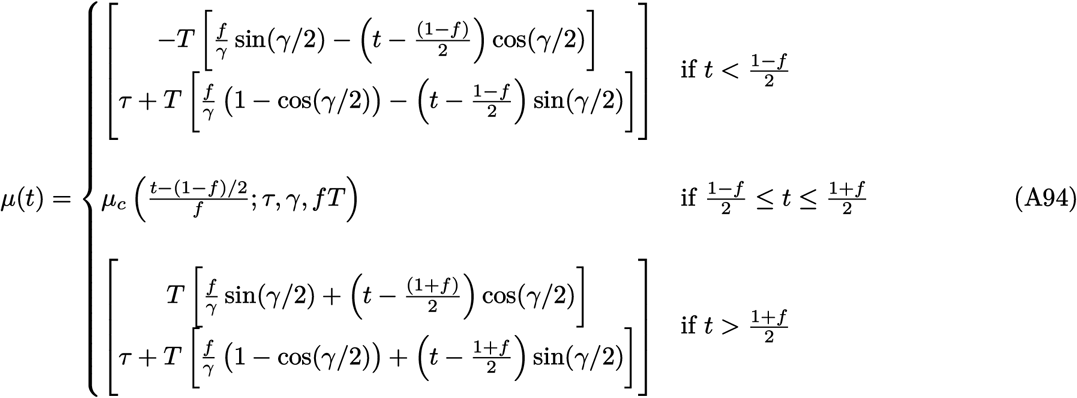

where *µ*_*c*_ is the *µ* mapping defined for the circular model, Equation A80.

For the experiment, the values of the parameters were chosen to guarantee that the circular section of this model would have the same center as a circular model (described above) with *γ* = (3*/*5)*π* and *τ* = 0, and that a relatively large fraction of the two models is in close proximity. The values are

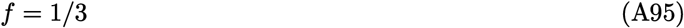

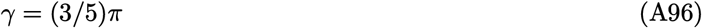

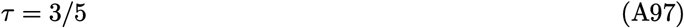

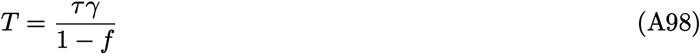

Closed-form expressions for all FIA terms can be derived for this model by a straightforward, if somewhat laborious, extension of those presented above for the circular-arc model. We do not report them here in the interest of brevity.

##### A.4.7 Point model

This model, used in the Dimensionality task variant, has no associated latent parameters (it is zero-dimensional). To cast it in the same language as the others, we can define it as

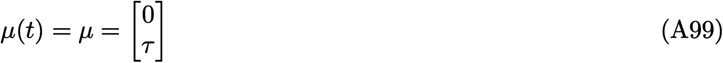

For the point model, the FIA (which is an approximation to a model’s log evidence) is replaced by the exact evidence, which simply coincides with the log likelihood. For notational consistency, we adopt the following values for the FIA terms:

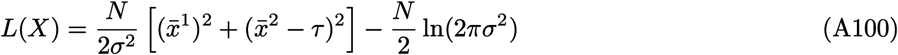

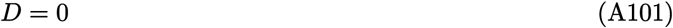

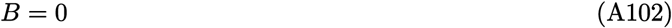

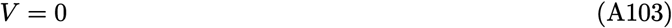

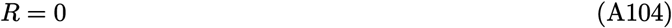

#### A.5 Numerical experiments with Artificial neural networks

##### A.5.1 Inputs

On each trial, our artificial neural network (“ANN”) takes as input: 1) two images, each depicting one candidate model’s location in the data space; and 2) a length-20 vector, containing the horizontal and vertical coordinates of the *N* = 10 data points. Each of the two images is provided as one RGB matrix of size (3*256*256). In data-space units (used for the model definitions in subsection A.4), each image extends from *x* = −4 to *x* = −4 and from *y* = −3.5 to *y* = 4.5, so that the center of the image (located in (0, 0.5)) is equidistant from the models in each model pair.

##### A.5.2 Training dataset

The training dataset consisted of 5000 model pairs. Each model pair was used for generating 50 trials. This approach led to a total of 250,000 trials in the entire dataset.

The random generation of model pairs was performed as follows (see subsection A.4 for the detailed mathematical definitions of each model and the precise meaning of the parameters controlling its shape). Each model pair could be of one of the four variants described in subsection A.3, chosen randomly with equal probabilities. Each model pair could be flipped vertically with probability 0.5. For the robustness variant, the separation of the model pair was 0.6 data-space units; for all other model pairs, the separation was 1 data-space unit. For the dimensionality variant, the length *T* (in data-space units) of the one-dimensional model was sampled uniformly from *U*(0.5, 5). For the boundary variant, the length of both model families were kept identical and sampled from *U*(0.5, 3). For the volume variant, the lengths of both models were sampled independently from *U*(0.5, 5); if their length difference was no greater than the task’s noise level *σ* = 1, then the length of one model was resampled from *U*(0.5, 5) until the length difference was greater than 1. For the robustness variant, the length of both model families was kept constant at (27*/*50) · *π*. The length proportion of the rounded model that was perfectly circular was *f* = 1*/*3, and both model families share the same curvature parameter *γ* sampled from *U*(1.5, 3). The model pairs were centered around the center of each input image.

Given a model pair, each trial was generated randomly as follows: 1) select one model randomly with equal probability; 2) sample a location along this model uniformly; and 3) using this location as the center of a 2D isotropic Gaussian and standard deviation of *σ* = 1 data-space units, sample *N* = 10 data points that were given to the network.

The training dataset was pre-shuffled randomly for training purposes. The input batch size was always 50 trials.

##### A.5.3 Test dataset

The test dataset consisted of 8 model pairs, each generating 15,000 trials. Thus, there was a total of 120,000 trials in the dataset.

The model pairs were as follows. For the Dimensionality task, the one-dimensional model had length (in data space units) 1. For the Boundary task, both model families had length 1. For the Volume task, one model had length 1, the other had 3. For the Robustness task, both model families had length *T* = (27*/*50) ·*π, f* = 1*/*3, and curvature parameter *γ* = (3*/*5)*π*. Each model pair was presented in the “upright” position (as per the definitions in subsection A.4) and in the vertically flipped position, for a total of 8 cases. The separation between model families and the generation of trials was identical to as in the training dataset.

##### A.5.4 Artificial neural network architecture

Our ANN had the following architecture (see Figure 4). Each of the two model input images was passed through the pretrained convolutional neural network VGG16, which had its parameters frozen during training. We replaced the fully connected layers at the end of VGG16 with our own structure of Linear-ReLU-BatchNorm1D layers and allowed the updating of weights in these and all subsequent layers. For each image input, the output of this image-processing module was a length-50 vector (model image representation).

In parallel, the length-20 vector of raw data point coordinates was fed through a permutation-invariant layer. This layer featured shared weights such that its outputs were not affected by the sequence of the N=10 data points in the length-20 vector input. This layer also outputted a length-20 vector, which was concatenated to the end of each of the length-50 vectors (the model image representations) along the preexisting dimension, producing two length-70 vectors.

Each length-70 vector was fed through Linear-ReLU-BatchNorm1D layers (identical weights used to process each vector). The resultant two length-50 output vectors were then concatenated together along the preexisting dimension, with the first input image’s representation in front.

The resultant length-100 vector was then fed through EquiLinear-ReLU-BatchNorm1D layers. The EquiLinear layers were permutation-equivariant layers of our design, again achieved by weight sharing. They ensure that if we concatenated the two length-50 output vectors in the opposite sequence, then their output, a length-2 vector, also had the same values but in opposite sequence. This length-2 vector was passed through a log softmax layer to produce the ANN’s final output, which was also a length-2 vector.

We also introduced a conditional variational encoder (CVAE) structure and used its output as part of the loss function (discussed later), to encourage model representations to preserve information about the data generation process. The details are described below.

We concatenated the length-20 raw data points vector (before passing input the permutation-invariant layers) to the end of each length-50 model image representation vector. The resultant two length-70 vectors (each corresponding to one model) were used as inputs for our CVAE (identical weights used to process each vector). The CVAE took each length-70 vector through its encoder structure to produce 10-dimensional vectors, which were used as parameters (*µ*_*CV AE*_, *σ*_*CV AE*_) for the Gaussian random generation of another 10-dimensional vector. The latter vector was again concatenated to the end of the length-50 model image representation vector responsible for its own generation, before being fed to the CVAE decoder, which mapped back to a 20-dimensional output vector reminiscent of data points. Hence, there were two 20-dimensional output vectors generated, each originating from one model.

##### A.5.5 Loss function

The loss function for each trial consisted of 2 parts: 1) the final output loss, and 2) the CVAE output loss. For the final output loss, we used Pytorch’s negative log likelihood loss function `NLLLoss()`, which computed the loss between the ANN’s length-2 output vector and the target label. For each trial’s CVAE output loss, we considered only the CVAE output associated with the correct model image/target label (hence one out of the two CVAE output vectors). The CVAE output loss was the sum of a MSE reconstruction loss (between the length-20 CVAE output vector and the length-20 raw data points vector) and a KL Divergence Loss (considering (*µ*_*CV AE*_, *σ*_*CV AE*_) used in the CVAE data generation process, using sum reduction). The total loss was the sum of the final output loss and the CVAE output loss.

##### A.5.6 Update rule

We used Pytorch’s Adam optimizer with learning rate 0.005, keeping all other arguments at their default values.

##### A.5.7 ANN predictions

To evaluate ANN task performance in a way that was comparable to human performance, we needed to specify how the ANN output, a length-2 log softmax vector, mapped onto a chosen candidate model. The mapping was as follows: we compared the two entries in the output vector and assumed that the ANN chose the candidate model associated with the larger entry.

#### A.6 Experimental data analysis

For both human and ANN experiments, we modeled behavior assuming that each observer samples from a posterior over models determined by a modified version of the FIA, where each term of the approximation is multiplied by a free parameter to be inferred, representing the sensitivity of the participant to that term.

Specifically, in our experimental scenario the theory of Bayesian model selection applies directly. Given two models ℳ_1_ and ℳ_2_, assuming a flat prior over models *p*(ℳ_1_) = *p*(ℳ_2_) = 1*/*2 and an uninformative prior (Jeffreys prior, see Balasubramanian (1997) and Jaynes (2003)) over the parameters of each model, when *N* is sufficiently large we can use the asymptotic expansion in Figure 1 and Equation A33 to write the log posterior ratio for ℳ_1_ over ℳ_2_ as

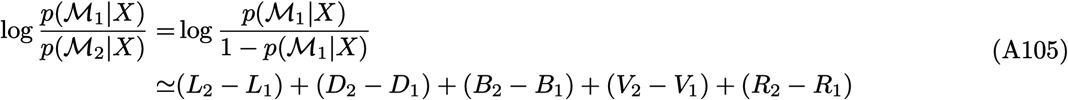

where *L*_*i*_, *D*_*i*_, etc represent the FIA terms for model *i*:

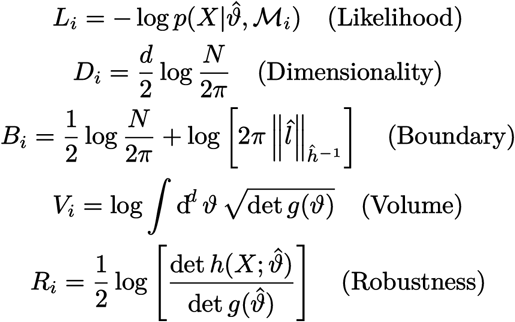

This expression suggests a simple normative model for participant behavior. Equation A105 determines the probability of reporting ℳ_1_ for an ideal Bayesian observer performing probability matching. We can then compare participant behavior to the normative prescription by allowing participants to have distinct sensitivities to the various terms of the FIA:

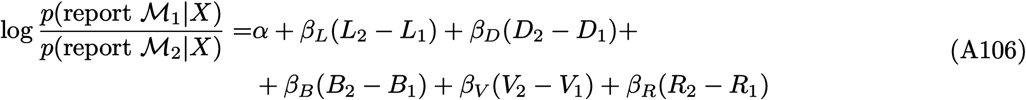

where *α* and *β* were free parameters: *α* captures any fixed bias, *β*_*L*_ the sensitivity to differences in maximum likelihood, *β*_*D*_ the sensitivity to differences in dimensionality, and so on.

We fitted the model expressed by Equation A106 to participant behavior using a hierarchical, Bayesian logistic-regression scheme:

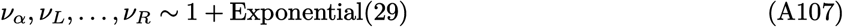

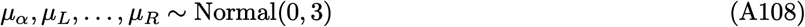

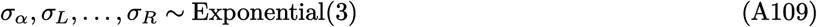

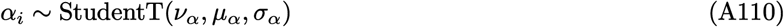

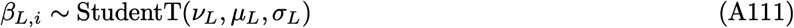

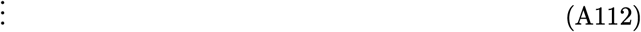

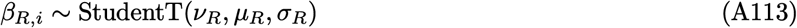

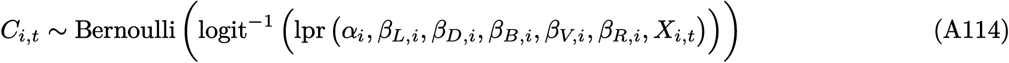

where *C*_*i,t*_ is the choice made by participant *i* on trial *t, X*_*i,t*_ is the sensory stimulus on that same trial, lpr is the log posterior ratio defined by Equation A106, *α*_*i*_ is the bias for participant *i, β*_*L,i*_ is the likelihood sensitivity of that same participant, and so on for the other sensitivity parameters. The bias and sensitivity parameters describing each participant are modeled as independent samples from a population-level Student-T probability distribution characterized by a certain shape (*ν*), location (*µ*), and scale (*σ*). The priors assumed over these population-level parameters are standard weakly informative priors (Gelman et al., 2014; Kruschke, 2015). The model was implemented in PyMC (Salvatier et al., 2016), and inference was performed by sampling from the posterior for the parameters given the experimental data {*C*_*i,t*_, *X*_*i,t*_*}* using the No-U-Turn Sampler algorithm (Betancourt, 2018; Hoffman & Gelman, 2014). Further technical details on the inference procedure can be found in subsubsection A.6.2.

##### Definition of relative sensitivity

Relative sensitivity for a certain feature was defined as the sensitivity for that feature divided by the relevant posterior mean for the likelihood sensitivity. For instance, for dimensionality:

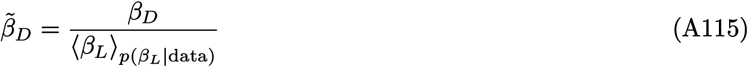

This formulation applies both at the participant level and at the population level.

Because each human participant performed only one task variant, not all sensitivities could be estimated for all participants. For instance, *β*_*D*_ entered the behavioral model (and therefore could be estimated) only for the participants that performed the Dimensionality task, where the alternative models had different dimensionality. The same holds with *β*_*V*_ and the Volume task, and *β*_*R*_ and the Robustness task. The boundary term entered the behavioral model for all task variants, although by design it took on a much broader range of values for the Boundary task. For consistency, for each sensitivity parameter, we reported its estimate only for those participants that performed the task variant designed to test it.

##### A.6.1 Lapse rates

We designed a variant of the behavioral model that accounts for lapses in participants’ responses (i.e., errors on easy trials). Specifically, we modified Equation A106 as follows:

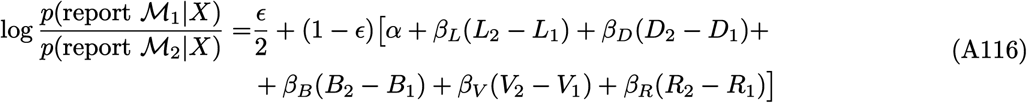

where *ϵ* ∈ [0, 1] is the lapse rate, representing the probability that a given response is completely random. For *ϵ* = 1 the responses are random on every trial, whereas for *ϵ* = 0 this model is equivalent to the original one in Equation A106.

To estimate *ϵ* from our experimental data jointly with all other parameters, we kept the same structure as in Equations A107–A114 and extended it by modeling the population level distribution of *ϵ* as a Beta distribution, parameterized by count parameters *a* and *b*. Following the recommendations by Gelman et al. (2014, section 5.3) and development team (2022, section 24.2), we specify hyperpriors in terms of the mean of the distribution *φ* = *a/*(*a* + *b*) and the total count *λ* = *a* + *b*:

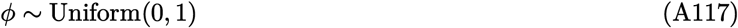

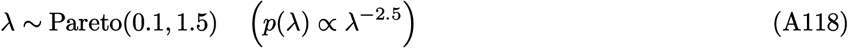

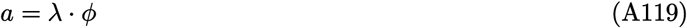

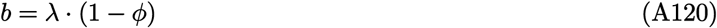

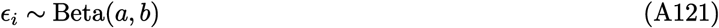

where *ϵ*_*i*_ is the lapse rate for participant *i*.

**Table A.1.** 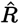 statistic and effective sample size (ESS) for 12 Markov Chain traces run as described in the text, for the fit to human data for the generative task. See Gelman et al. (2014, sections 11.4–11.5) and Vehtari et al. (2020) for in-depth discussion of chain quality diagnostics. Briefly, 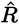 depends on the relationship between the variance of the draws estimated within and between contiguous draw sequences. 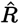 is close to 1 when the chains have successfully converged. The effective sample size estimates how many independent samples one would need to extract the same amount of information as that contained in the (correlated) MCMC draws.

| Parameter | ESS | $\hat{R}$ |
| --- | --- | --- |
| $\mu_\alpha$ | 52148 | 1.000 |
| $\mu_L$ | 19254 | 1.000 |
| $\mu_D$ | 27384 | 1.000 |
| $\mu_B$ | 32434 | 1.000 |
| $\mu_V$ | 64614 | 1.000 |
| $\mu_R$ | 112118 | 1.000 |

##### A.6.2 Technical details of the inference procedure

Posterior sampling was performed with PyMC (Salvatier et al., 2016) version 4.2.0, using the NUTS Hamiltonian Monte Carlo algorithm (Hoffman & Gelman, 2014). Target acceptance probability was set to 0.9 for the human data (both generative and maximum-likelihood task), to 0.8 for the generative task for ANNs, and 0.99 for the maximum-likelihood task for ANNs. The posterior distributions were built by sampling 12 independent Markov chains for 10000 draws each. No divergence occurred in any of the chains. Effective sample size and 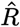 diagnostics for some of the key parameters are given in Table A.1.

##### A.6.3 Reporting of posterior distributions for inferred parameters

The posterior distributions reported in all figures are Kernel Density Estimates with bandwidth chosen according to Scott’s rule (Pedregosa et al., 2011).

#### A.7 Fitting the NIN observer to participant data

We fit the NIN model to our human behavioral data by maximum likelihood, on a single-participant basis and marginalizing numerically over the effect of sensory noise. In particular, say that 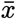 is the centroid of the dot cloud. Recall that the NIN model involves a latent (internal) sensory noise process that corrupts the sensory data by adding noise *η* ∼ *N*(0, *ρ*):

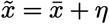

Then, the model observer computes a Bayesian posterior 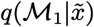 by integrating locally around the maximum-likelihood point 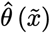. The range of the local integration is controlled by an integration parameter *b*, so we write

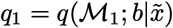

Finally, choice noise is implemented through a softmax transform:

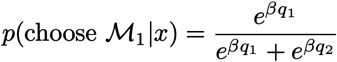

where *β* = 1*/T* is the “inverse temperature” parameter controlling the intensity of the choice noise. Inference is based on calculating the (log)likelihood of the model (parameterized by *ρ, b* and *β*), obtained by marginalizing over the sensory noise *η*:

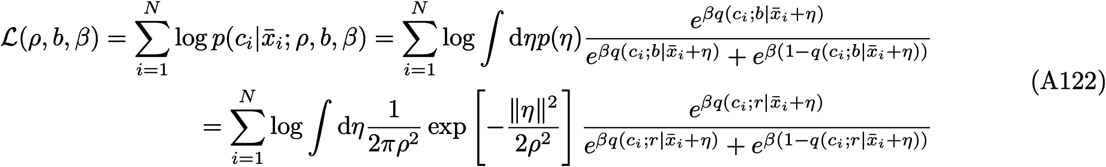

We approximate the integral in Equation A122 using a randomized quasi-Monte Carlo (QMC) approach (Owen, 2023):

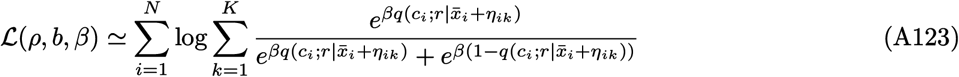

where 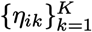, for any *i*, is a bivariate normal QMC set of points generated from a scrambled Sobol’ sequence with = 1024 using the `scipy.stats.qmc` module (Roy et al., 2023), centered in (0, 0) and with variance *ρ* . We used an independent set of QMC points for each participant and each trial, but we kept this set fixed throughout the optimization (that is, we did not resample the QMC set at each iteration of the optimization procedure) to avoid introducing stochasticity in the loss function and to help with convergence.

To fit the model, we maximised the likelihood Equation A123 using Differential Evolution (Storn & Price, 1997) as implemented in the `scipy.optimize` module with a candidate population of size 64. The model parameters were constrained to the following ranges: *ρ* ∈ (0, 2], *b* ∈ [0, 1], *β* ∈ [0, 10].

## Appendix B

### Supplementary information

#### B.1 Numerical comparison of the extended FIA vs exact Bayes

**Figure B.1.**
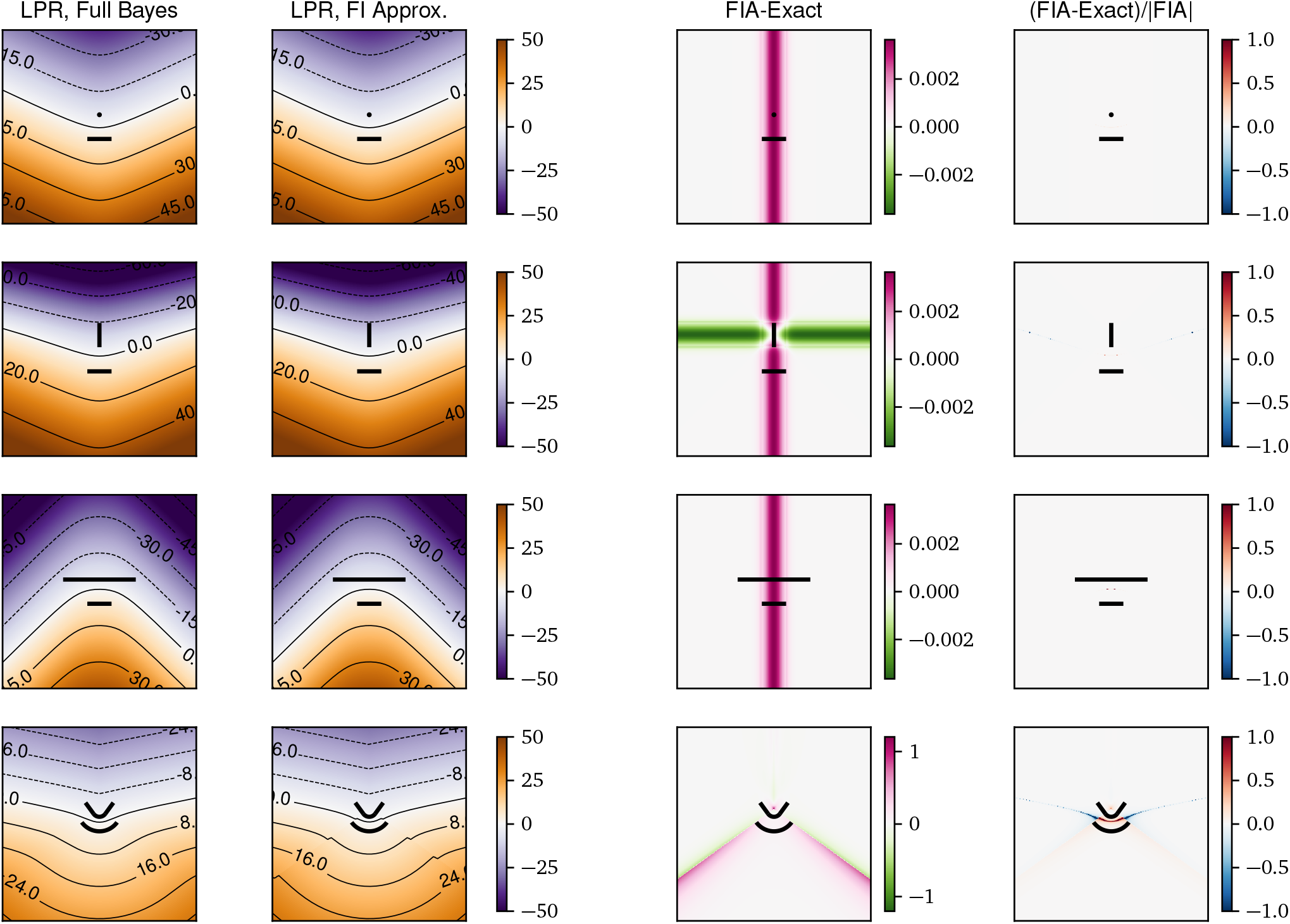
Comparison of the full Bayesian and FIA computation of the log posterior ratio (LPR) for the model pairs used in our psychophysical tasks (N = 10). Each row corresponds to one task variant (from top to bottom, “dimensionality”, “boundary”, “volume”, “robustness”). First column from the left: full Bayesian LPR, computed by numerical integration. Second column: LPR computed with the FIA. Third column: difference between FIA and exact LPR. Fourth column: relative difference (difference divided by the absolute value of the FIA LPR). Adapted from Piasini et al. (2021a).

Figure B.1 shows that the FIA computed with the expressions given in this document provides a very good approximation to the exact Bayesian log posterior ratio (LPR) for the model pairs used in the psychophysics experiments, and for the chosen sample size (*N* = 10). As highlighted in the panels in the rightmost column, the discrepancies between the exact and the approximated LPR are generally small in relative terms, and therefore are not very important for the purpose of model fitting and interpretation. Note that here, as well as for the results in the main text, the *B* term in the FIA is computed using Equation A34 rather than Equation A38 to avoid infinities (that for finite *N* can arise when the likelihood gradient is very small) and discontinuities (that for finite *N* can arise on the interior of the manifold, in proximity to the boundary, where the value of *B* goes from zero when 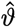 is in the interior to log(2) when 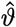 is exactly on the boundary).

Even though overall the agreement between the approximation is good, it is interesting to look more closely at where the approximation is poorest. The task type for which the discrepancies are the largest (both in absolute and relative terms) is the “robustness” type (fourth row in Figure B.1). This discrepancy arises because the FIA hypotheses are not fully satisfied everywhere for one of the models. Specifically, the models in that task variant are a circular arc (the bottom model in Figure B.1, third row) and a smaller circular arc, concentric with the first, with a straight segment attached to either side (the top model). The log-likelihood function for this second model is smooth only to first order, but its second derivative (and therefore its Fisher Information and its observed Fisher Information) is not continuous at the points where the circular arc is joined with the straight segments, locally breaking hypothesis number 3 in subsubsection A.2.1. Geometrically, this discontinuity is analogous to saying that the curvature of the manifold changes abruptly at the joints. It is likely that the FIA for a model with a smoother transition between the circular arc and the straight arms would have been even closer to the exact value for all points on the 2D plane (the data space). More generally, this line of reasoning suggests that it would be interesting to investigate the features of a model that affect the quality of the Fisher Information Approximation.

#### B.2 Posterior predictive checks

**Figure B.2.**
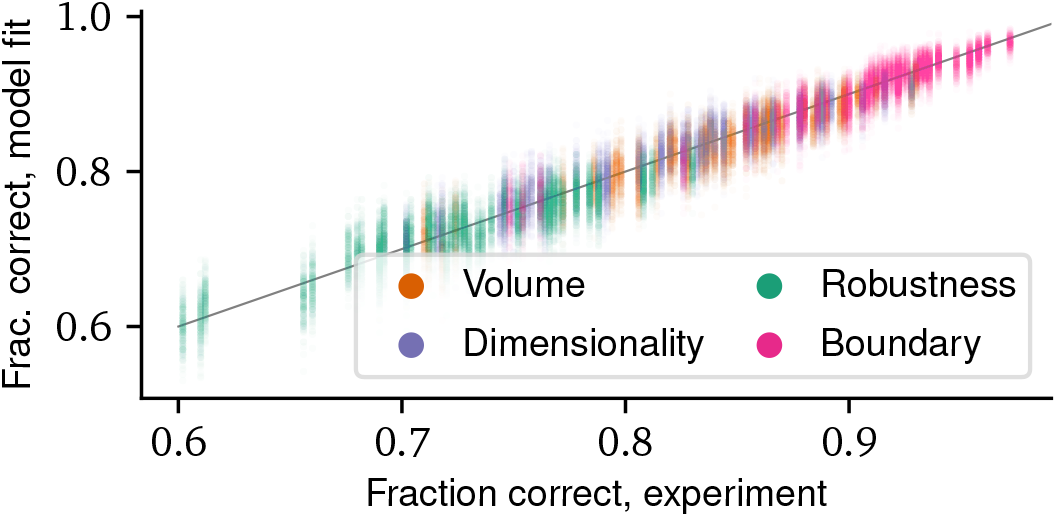
Posterior predictive check for human performance on the generative task. We sampled 240 samples from the posterior over model parameters by thinning the MCMC chains used for model inference. For each of these samples, we ran a simulation of the experiment using the actual stimuli shown to the participants, and we recorded the resulting performance of all 202 simulated participants. This procedure yielded 240 samples of the joint posterior-predictive distribution of task performance over all participants. To visualize this distribution, for each participant we plotted a cloud of 240 dots, where the y coordinate of each dot is the simulated performance of that participant in one of the simulations, and the x coordinate is the true performance of that participant in the experiment plus a small random jitter (for ease of visualization). The gray line is the identity, showing that our inference procedure captures well the behavioral patterns in the experimental data. Colors indicate different task types, as indicated.

We performed a simple posterior predictive check (Kruschke, 2015) to ensure that the Bayesian hierarchical model described in the text captures the main pattern of behavior across our participants. In Figure B.2, the behavioral performance of the participants is compared with its posterior predictive distribution under the model, for the case of the human participants in the generative task. As can be seen from the figure, the performance of each participant is correctly captured by the model, across systematic differences between task types (with participants performing better in the boundary task variant than the robustness task variant, for instance) as well as individual differences between participants that performed the same task variant.

#### B.3 Details on raw estimated sensitivities

Table B.3 reports the posterior mean and standard deviation of the population-level parameters entering the regression (Equation A106). Note that these are the raw parameters, not their normalized counterparts relative to the likelihood sensitivity as reported in the rest of the paper.

#### B.4 Uncertainty in participant-level sensitivities

Figure B.4 illustrates the uncertainty in the estimate for the relative sensitivity of each participant. This uncertainty is typically small compared to between-participant variability of the sensitivity, which is therefore not a trivial consequence of the noise in the sensitivity estimation for individual participants.

**Figure B.4.**
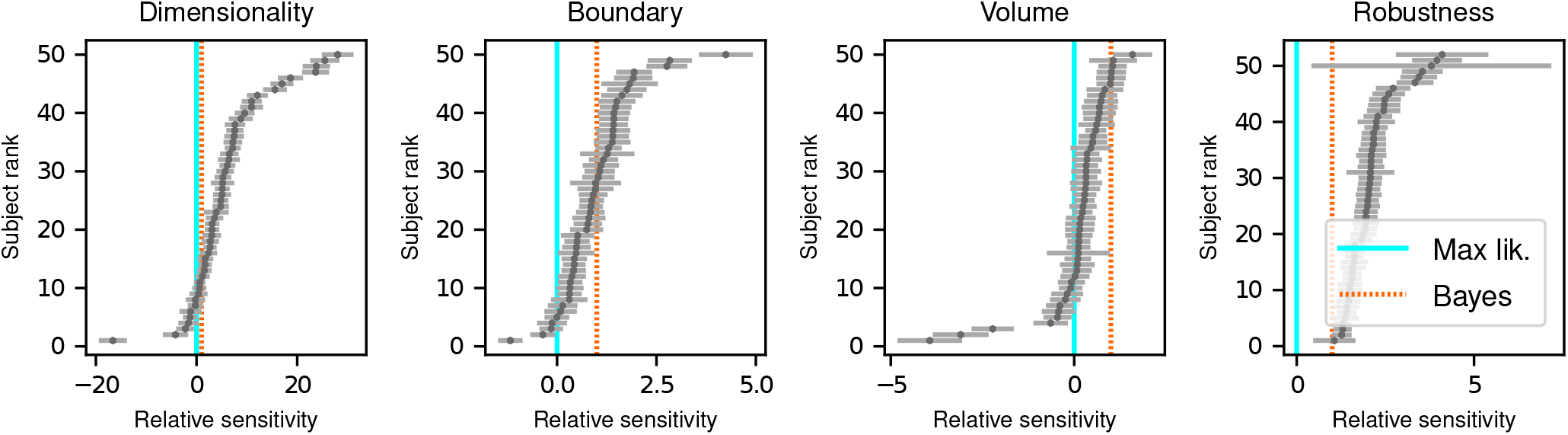
Participant-level relative sensitivities to the geometric features that determine model complexity. Dots with error bars: posterior mean ± standard deviation of the relative sensitivity (the dots are the same as in Figure 3c). For ease of visualization, participants are ranked based on their posterior mean.

**Table B.3.** Posterior mean±standard deviation for population-level parameters. See Equation A106 to Equation A114 for the precise definition of each parameter and its role in the hierarchical model of behavior.

| Parameter | Generative task |  | Max likelihood task |  |
| --- | --- | --- | --- | --- |
|  | Humans | ANNs | Humans | ANNs |
| $\mu_\alpha$ (up/down bias) | $0.107 \pm 0.023$ | $-0.242 \pm 0.151$ | $0.056 \pm 0.024$ | $0.010 \pm 0.07$ |
| $\mu_L$ (likelihood) | $0.461 \pm 0.012$ | $6.529 \pm 0.188$ | $0.561 \pm 0.018$ | $7.966 \pm 0.401$ |
| $\mu_D$ (dimensionality) | $2.150 \pm 0.445$ | $8.484 \pm 0.653$ | $1.285 \pm 0.231$ | $-0.030 \pm 0.486$ |
| $\mu_B$ (boundary) | $0.518 \pm 0.045$ | $6.286 \pm 0.186$ | $0.499 \pm 0.058$ | $0.883 \pm 0.121$ |
| $\mu_V$ (volume) | $0.108 \pm 0.057$ | $6.089 \pm 0.204$ | $0.105 \pm 0.044$ | $0.128 \pm 0.196$ |
| $\mu_R$ (robustness) | $1.018 \pm 0.056$ | $4.882 \pm 0.417$ | $1.276 \pm 0.085$ | $0.356 \pm 0.281$ |

#### B.5 Lapse-rate analysis

To ensure that our findings were not sensitive to possible lapses in attention by the participants, we used the model variant described in Section A.6.1 to estimate a lapse rate for each participant simultaneously with the sensitivity parameters (Figure B.5). There was a substantial spread of lapse rates in the range 0–0.2 but no clear relationship between lapse rates and sensitivity. The sensitivity parameters estimated with this extended model were qualitatively compatible with those presented everywhere else in the text.

#### B.6 Outcome of significance tests specified in the preregistration documents

##### B.6.1 Formal comparison between ideal observers

We compared the Bayesian hierarchical model described in section A.6 to a simpler model, where participants were assumed to be sensitive only to likelihood differences, or in other words to choose ℳ_1_ over ℳ_2_ based only on which model was on average closer to the dot cloud constituting the stimulus on a given trial. Mathematically, this “likelihood-only” model was equivalent to fixing all *β* parameters to zero except for *β*_*L*_ in the model described in section A.6. All other details of the model were the same, and in particular the model still had a hierarchical structure with adaptive shrinkage (the participant-level parameters *α* and *β*_*L*_ were modeled as coming from Student T distributions controlled by population-level parameters). We compared the full model and the likelihood-only model on our human behavior data using WAIC (Gelman et al., 2014). This comparison indicated strong evidence in favor of the full model for both the generative task (Table B.6) and the maximum-likelihood task (Table B.7).

**Figure B.5.**
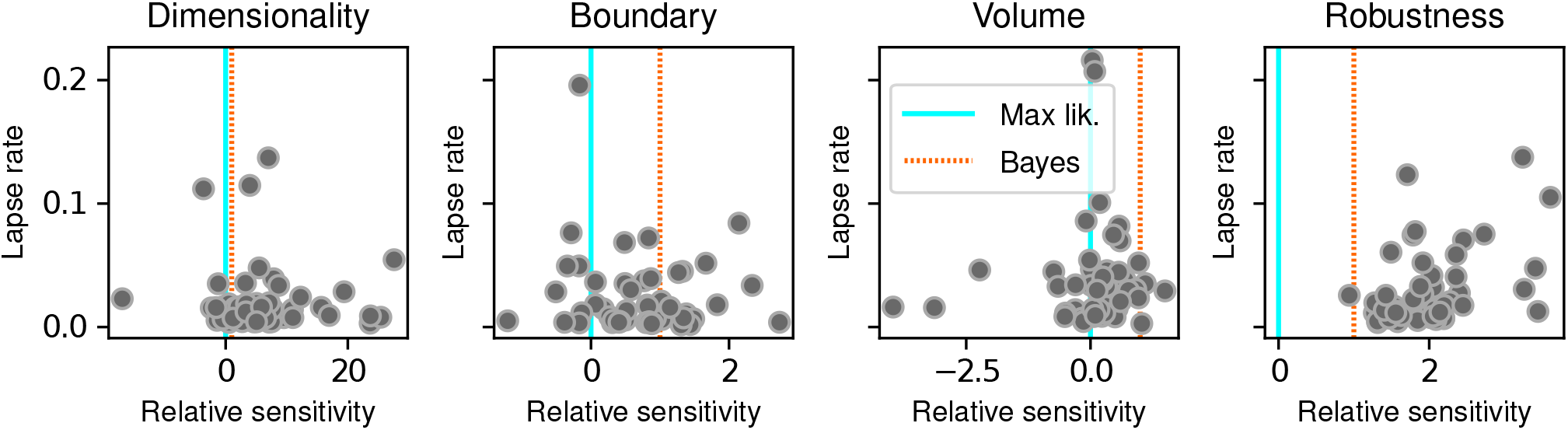
Lapse rate versus relative sensitivity to complexity across participants. Each dot gives the posterior mean estimate of the relative sensitivity to one of the features that determine model complexity (abscissa) and the posterior mean estimate of the lapse rate, as defined in Section A.6.1.

**Table B.6.** WAIC comparison of the full model and the likelihood-only model for human performance on the generative task, reported in the standard format used by McElreath (2016, section 6.4.2). WAIC is the value of the criterion (log-score scale, where higher is better), pWAIC is the estimated effective number of parameters, dWAIC is the difference between the WAIC of the given model and the highest-ranked one, SE is the standard error of the WAIC estimate, and dSE is the standard error of the difference in WAIC. These estimates were produced with the compare function provided by ArviZ (Kumar et al., 2019), using 12 MCMC chains with 10000 samples each for each model (in total, 120,000 samples for each model).

| Model | Rank | WAIC | pWAIC | dWAIC | SE | dSE |
| --- | --- | --- | --- | --- | --- | --- |
| Full | 0 | -34824.6 | 641.963 | 0 | 183.981 | 0 |
| Likelihood only | 1 | -37524.9 | 370.340 | 2700.4 | 183.923 | 69.3817 |

##### B.6.2 Other statistical tests

As described in the preregistration documents (Piasini et al., 2020, 2021b, 2022), in this work we have emphasized parameter estimation and information criteria-based model comparison over null hypothesis significance testing (see for instance McElreath (2016), and Kruschke (2015) for a discussion and comparison of these ideas). However, for completeness, we report in Table B.8 (1) the comparison between the Regions of Practical Equivalence (ROPE, Kruschke (2015)) and the 95% highest-density interval (HDI) for each population-level parameter, and (2) the “probability of direction” (Makowski, Ben-Shachar, Chen, & Lüdecke, 2019; Makowski, Ben-Shachar, & Lüdecke, 2019) for the same parameters (see below for more details on these methods). The ROPE-HDI tests highlight that the null value of zero sensitivity is not credible (rejected) for *L, D* and *R*, and neither rejected not accepted for *V* . The probability of direction is high for all parameters, including *V*, which has *PD* = 0.97 for the generative task and *PD* = 0.99 for the maximum-likelihood task. Overall, these analyses point to a significant sensitivity for all terms of the FIA in both experiments (generative and maximum-likelihood), with *V* having a more moderate effect size than the other terms.

###### Technical details on the ROPE-HDI comparison and on the Probability of Direction for sensitivity parameters

Briefly, the ROPE for a parameter is the range around a null value for that parameter such that variations within this range would imply only a “negligible change” in the behavior of the model, if all other parameters were held at their null values. The HDI is the smallest interval that contains a certain probability mass for the posterior of that parameter. The ROPE-HDI comparison is based on the idea that if the bulk of the posterior distribution for that parameter (represented by the HDI) falls outside the ROPE, then the null value for that parameter can be considered not credible (rejected). On the other hand, if the bulk of the posterior for the parameter falls within the ROPE, the null value can be considered credible (accepted). Finally, if the posterior distribution has a partial overlap with the ROPE (neither mostly contained within it, nor mostly falling outside of it), then the test is inconclusive. Note that, just like frequentist null hypothesis significance testing procedures and unlike the information criterion approach used above, this method depends on some arbitrary assumptions, namely the definition of the ROPE and the probability to use in computing the HDI.

**Table B.7.** Same as Table B.6, for the maximum-likelihood task, where participants were asked to report the model that was closest to the data.

| Model | Rank | WAIC | pWAIC | dWAIC | SE | dSE |
| --- | --- | --- | --- | --- | --- | --- |
| Full | 0 | -31022.8 | 638.926 | 0 | 184.912 | 0 |
| Likelihood only | 1 | -33155.1 | 374.023 | 2132.28 | 186.667 | 63.1851 |

**Table B.8.**
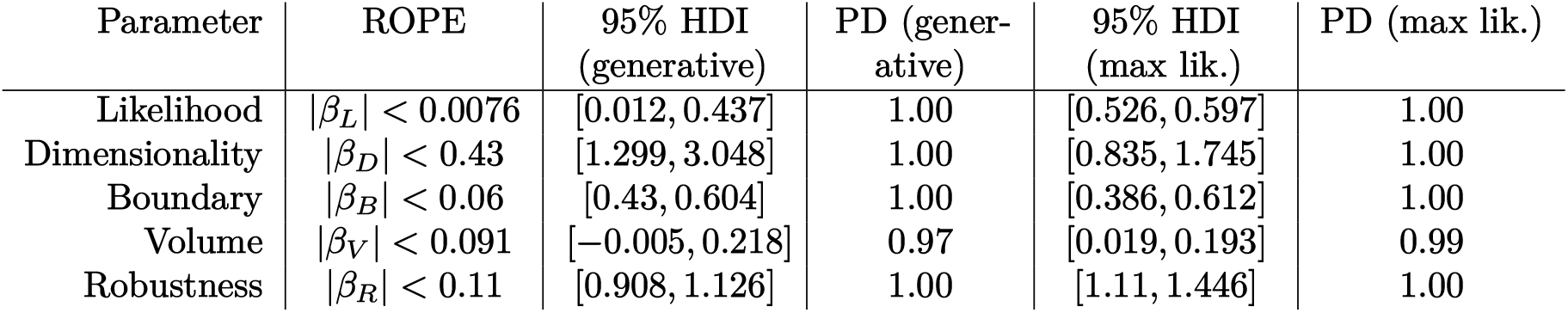
HDI vs ROPE comparison and Probability of Direction (PD) for the population-level parameters in the human experiments. See Supplementary Information section B.6.2 and Kruschke (2015) for an explanation of the ROPE-HDI comparison, and Makowski, Ben-Shachar, Chen, and Lüdecke (2019) and Makowski, Ben-Shachar, and Lüdecke (2019) for more details on the probability of direction metric. Note that the ROPE and HDI definitions were preregistered (Piasini et al., 2020, 2021b, 2022).

In practice (for details, see our preregistration documents (Piasini et al., 2020, 2021b, 2022)), here we define, conventionally, the HDI as the smallest interval that contains 95% of the posterior. The ROPE is computed as follows. We start by defining a “negligible change” over the probability of the choice variable over the “main range” [*µ*_*x*_ − 2*σ*_*x*_, *µ*_*x*_ + 2*σ*_*x*_] of one of the predictors in our model (*L, D, B, V*, or *R*). In other word, pick an interval of probabilities [*π*_0_ − *δ, π*_0_ + *δ*] such that if the probability stays within [*π*_0_ −*δ, π*_0_ + *δ*] when *x* varies over its typical range, then the probability is not meaningfully affected by *x*. Mathematically, if the probability of choosing one of the alternatives in the task is *π* and the log-odds is logit(*π*) = log(*π/*(1 − *π*)) =, then in a logistic regression setting

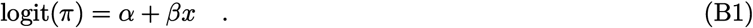

If *π*_0_ = logit^−1^(*α*), then the ROPE for *β* is defined as

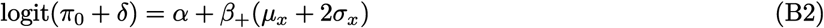

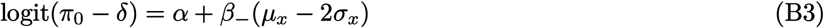

so that

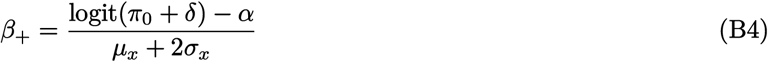

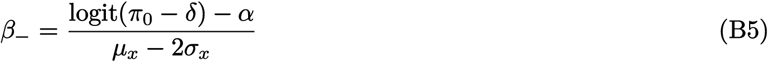

In our case, assuming a negligible influence of the up/down bias (*α* in Equation A106), *π*_0_ = 0.5, and therefore we can assume *α* = 0. The definition of the ROPE further depends on the arbitrary choice of *δ*, and on the values of *µ*_*x*_ and *σ*_*x*_. We choose *δ* = 0.025, and we estimated *µ*_*x*_ and *σ*_*x*_ by generating 25,000 experimental trials per task type (Dimensionality, Boundary, Volume, Robustness) and computing the empirical average and standard deviation of the predictors over that trial set. These numbers were all fixed at preregistration time (Piasini et al., 2020, 2021b, 2022).

#### B.7 Further details and results on the Noise-Integration-Noise observer

**Figure B.9.**
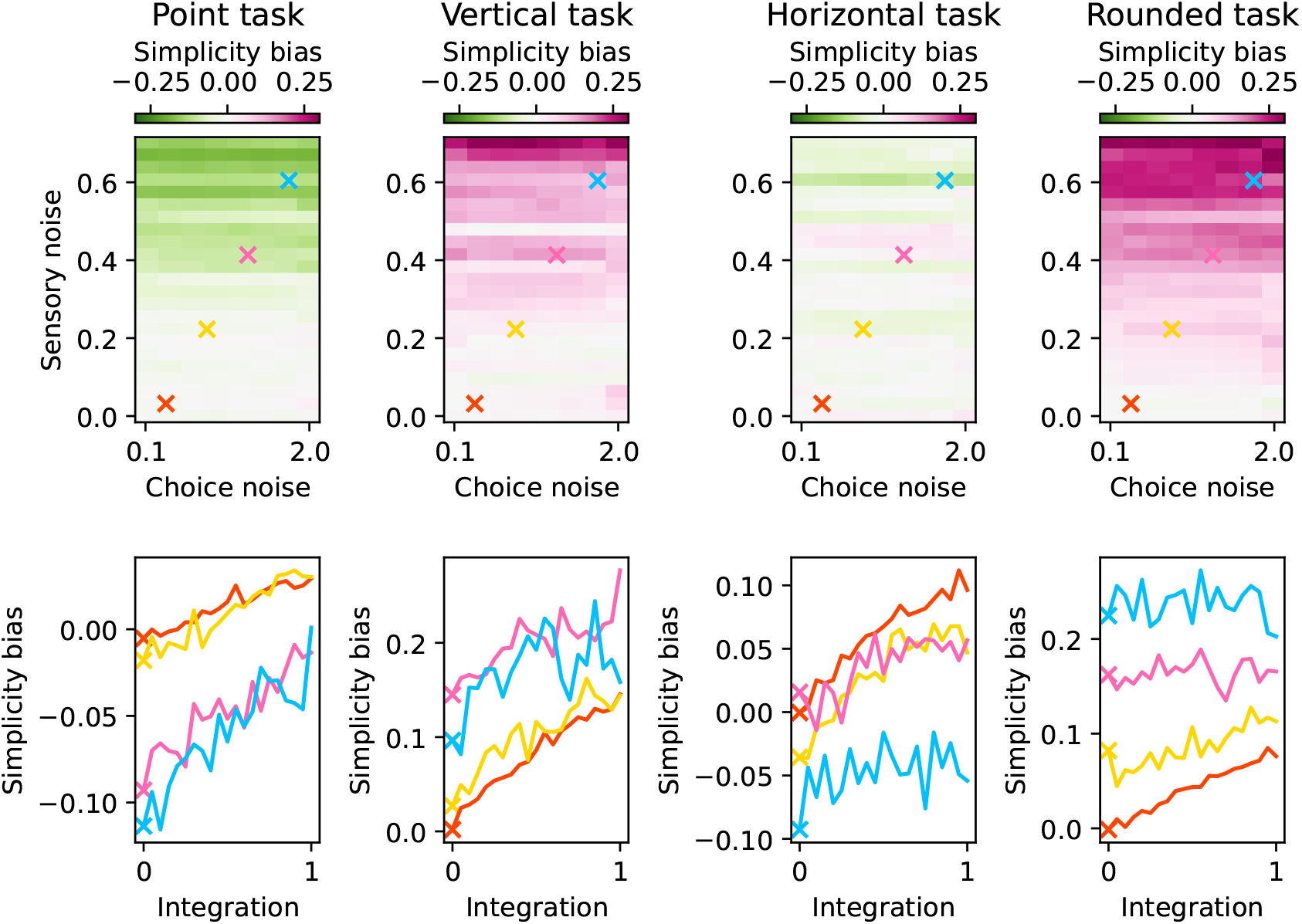
Model-free estimate of simplicity bias in the Noise-Integration-Noise (NIN) observer, as a function of the observer’s parameters, for each of the four task types (Dimensionality, Boundary, Volume, and Robustness; different columns show results for different task types). The example in Figure 2b in the main text corresponds to the Dimensionality task type. Top: simplicity bias as a function of sensory noise ρ and choice noise T, when the integration parameter b is set to 0, meaning that the observer does not integrate over latent causes (see section A.1). Note that the grid of choice noise values tested in the figure is not equally spaced; the values of T shown here are [0.10, 0.50, 0.56, 0.63, 0.71, 0.83, 1.00, 2.00], which correspond to the following values for the inverse temperature 1/T: [10, 2.0, 1.8, 1.6, 1.4, 1.2, 1.0, 0.5]. Bottom: simplicity bias as a function of the integration parameter b, with sensory noise and choice noise fixed to the values indicated in the top panels with the colored crosses. Note how integration is the only parameter that is associated with consistent changes in the simplicity bias for all task types (increasing integration increases the simplicity bias). Sensory noise has inconsistent effects across task types, and choice noise does not affect the simplicity bias.

##### B.7.1 Estimation of the simplicity bias in the NIN observer

To estimate the magnitude and source(s) of simplicity biases exhibited by the Noise-Integration-Noise (NIN) observer, for a given configuration of the observer we started by simulating 10000 trials for each task type. Using this simulated data, we first quantified the bias in a model-free way, following the procedure outlined in A.1.1. This analysis showed that integration, and not sensory or choice noise, is the only parameter that is associated with consistent changes in the simplicity bias for all task types (Figure B.9).

We then measured the sensitivity of the NIN observer to the terms of the FIA expansion. To do so, we fitted to the simulated data a simplified version of the model we used for humans and artificial neural networks, described in detail in A.6. Briefly, the log odds for the observer to choose model ℳ_1_ over ℳ_2_ was taken to be similar to Equation A106:

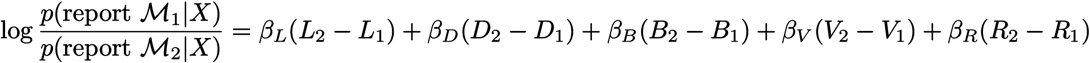

where *L*_*i*_, *D*_*i*_, *B*_*i*_, *V*_*i*_ and *R*_*i*_ are the FIA expansion terms for model M_*i*_, and *β*_*k*_ are parameters to be fitted, with, for instance, *β*_*D*_ representing the sensitivity of the observer to model dimensionality. We fit this expression to the behavior of the observer in the 10000 simulated trials using a regularized logistic regression:

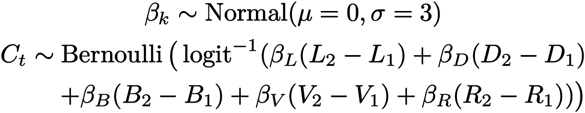

where *C*_*t*_ is the choice of the observer on model *t*, with *C*_*t*_ = 1 indicating that the observer chooses to report model ℳ_1_ on that trial. For each NIN configuration tested, we report in Figure B.10 the maximum a posteriori (MAP) fit of the *β* values, obtained using the `find_MAP` function in PyMC (Salvatier et al., 2016). Our results were broadly consistent to those in Figure B.9, showing that integration was the only observer parameter with a consistent effect on FIA term sensitivity (Figure B.10, top and middle). Finally, we normalized the FIA term sensitivity by the likelihood sensitivity, in the same way as we did in the main text for human and artificial neural network observers. This analysis showed an even stronger qualitative match between the model-free and the FIA-based quantification of the simplicity bias (compare Figure B.9 and Figure B.10, bottom).

##### B.7.2 Human simplicity preference is compatible with integration over latent causes

Fits of the Noise-Integration-Noise (NIN) model indicated that the behavior of most of our human participants was consistent with integrating over latent causes. Specifically, for thefor the Robustness and Dimensionality versions of the generative task most participants had their “integration indices” *b* pinned at 1, which is the maximum possible value and indicated integration over the full statistical manifolds (Figure B.11b). For the Volume and Boundary tasks, some participants failed to integrate (*b* = 0) but the others showed a range of propensities toward integration. These results are consistent with the sensitivities extracted with the FIA analysis (Table B.3), which showed that participants were generally more strongly sensitive to dimensionality and robustness than volume and boundary.

The distribution of best-fitting values of the sensory noise parameter *ρ* (Figure B.11a) exhibited two components: 1) a spike at zero, indicating a number of participants with negligible sensory noise; and 2) a separate, broader peak. This peak was centered around an “efficent” value, as we will now argue. As a reminder, on each trial of the experiment we showed a cloud of *N* = 10 dots to the participants, with each dot being sampled from a 2D Gaussian distribution with isotropic variance and standard deviation *σ* = 1. For our task, the location of the cloud centroid 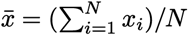 is a sufficient statistic, in the sense that it is enough to compute the posterior probabilities for our behavioral models (both FIA and NIN). The standard deviation of the sampling distribution for 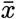 (its standard error) therefore captures the intrinsic reliability of the raw visual information provided to the participants, before it is corrupted by the sensory noise *η* ∼ N (0, *ρ*). From the definition of 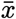, its standard error is simply 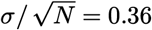. The variance of the perceived centroid location 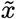 is then *σ*^2^*/N* + *ρ*^2^, which is dominated by *ρ*^2^ when 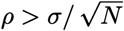 and by 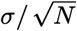 when 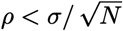. Therefore, 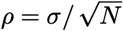 is the “efficient” value of *ρ*, corresponding to the maximum spatial resolution beyond which it is not convenient to encode the location of the centroid due to diminishing returns. This value is reported in Figure B.11a as an arrow, showing that the estimated values of *ρ* for the participants with non-negligible sensory noise are centered around this value.

The best-fitting inverse-temperature parameters have a distribution with two components (Figure B.11c): 1) a bulk below 1, and 2) a tail with values that go far above 1. As a reminder, for this parameter, 0 corresponds to making random choices, 1 corresponds to sampling from the posterior over models (computed by integrating in a neighborhood of the max likelihood point whose radius is determined by the “integration” parameter *b*), and infinity corresponds to selecting deterministically the model with the largest posterior. Also recall from A.7 that the inverse temperature *β* is bounded in the fitting procedure to the interval [0, 10], so the participants with *β* = 10 could possibly be fit even better by larger values. However, in practice this constraint does not make a large difference with respect to the behavior of the fitted model, as *β* = 10 can be already considered close enough to a deterministic choice regime. The peak of the inverse-temperature distribution is very close to the mean posterior estimate for the (population-level) sensitivity to likelihood in our FIA model (which is 0.46 in the generative task, plotted as a dashed cyan line in Figure B.11c). This result is consistent with the idea that the likelihood sensitivity captures an overall scaling of the slope of the psychometric function, and further supports our choice to focus on normalized sensitivities in the main text.

Applying the same analysis to the data from the maximum-likelihood task gives broadly consistent results, with the exception that the inferred degree of integration is less peaked around 1 and much more distributed over a continuum between 0 and 1 for all tasks (Figure B.12). This result implies that the participants still performed integration under these conditions, albeit to a lesser extent than for the generative task, corroborating the results of the FIA analysis in Figure 5 in the main text.

##### B.7.3 Model comparison between NIN and FIA

We checked whether our more refined, theory-driven behavioral model based on the FIA explained participant behavior better than the more elementary NIN model. To do so in a fair way, we re-fit the FIA model on a individual-participant basis (that is, we removed the hierarchical structure) using maximum likelihood, with the same optimization algorithm described for the NIN model in A.7. This procedure allowed us to compute an Akaike Information Criterion for each model on each participant. Figure B.11b shows that the vast majority of participants (182 out of 202) were better described by the FIA model.

**Figure B.10.**
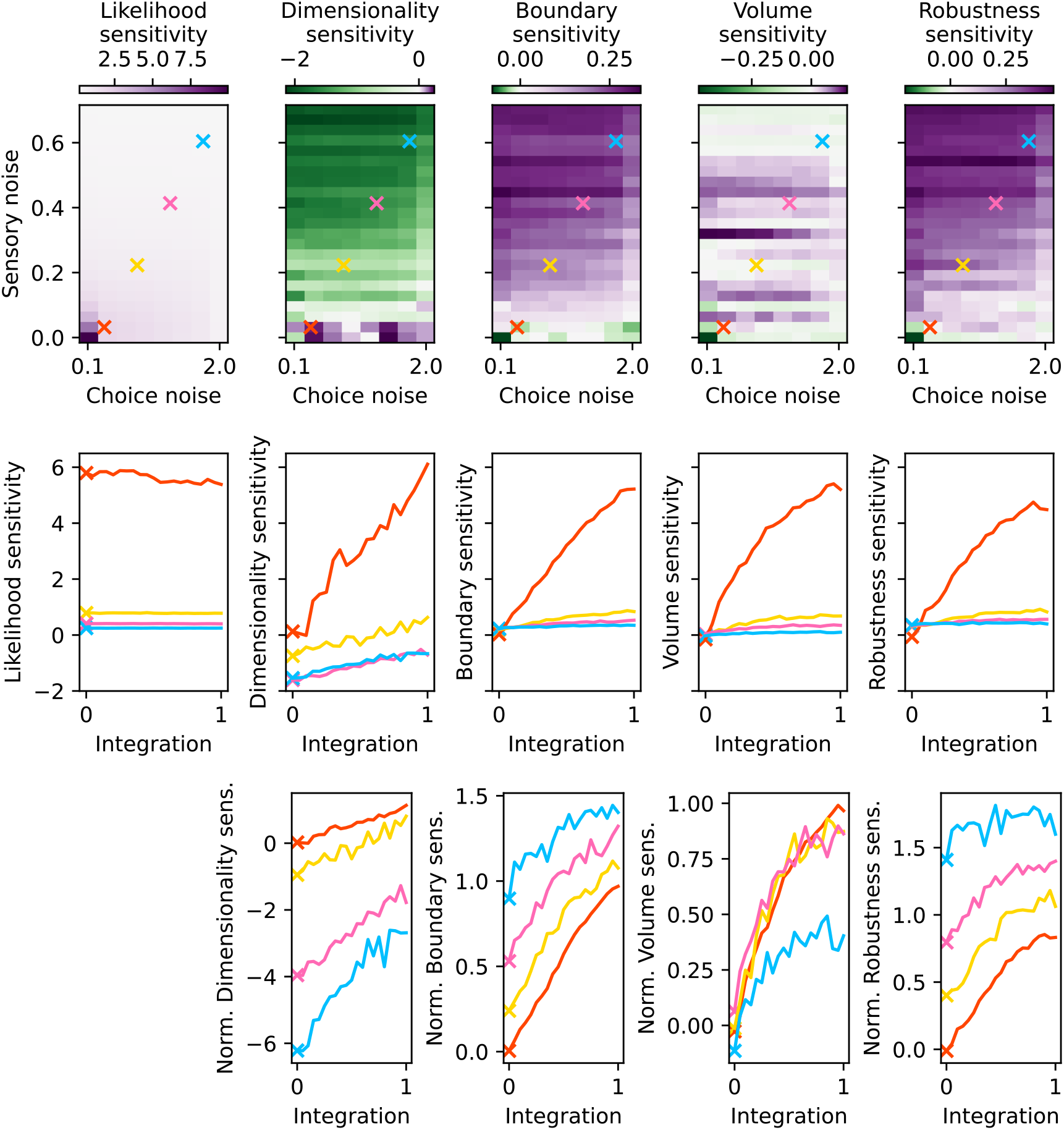
Sensitivity to model likelihood and to the geometric features that characterize model complexity, for the Noise-Integration-Noise observer, as a function of the parameters of the observer. The parameter values tested are the same as in Figure B.9. Top: dependence of the sensitivities on the sensory noise σ and the choice noise T, when the integration parameter b is fixed to zero (meaning that the observer does not integrate over latent causes). Middle: dependence on the sensitivities on integration, when sensory and choice noise are fixed to the values indicated by the colored crosses in the top panels. Bottom: same as middle, but for the normalized sensitivities, obtained by dividing the raw sensitivities by the likelihood sensitivity. Note that, reflecting the results in Figure B.9, the parameter controlling integration (x axis on each individual subplot) is the only one that has a consistent effect on the sensitivity to all features, generally increasing it. Note also the qualitative match between the bottom panels here and those in Figure B.9. The agreement with the data presented here and that in Figure B.9 further confirms that our theory-driven approach captures the intuitive notion of simplicity bias in this task.

**Figure B.11.**
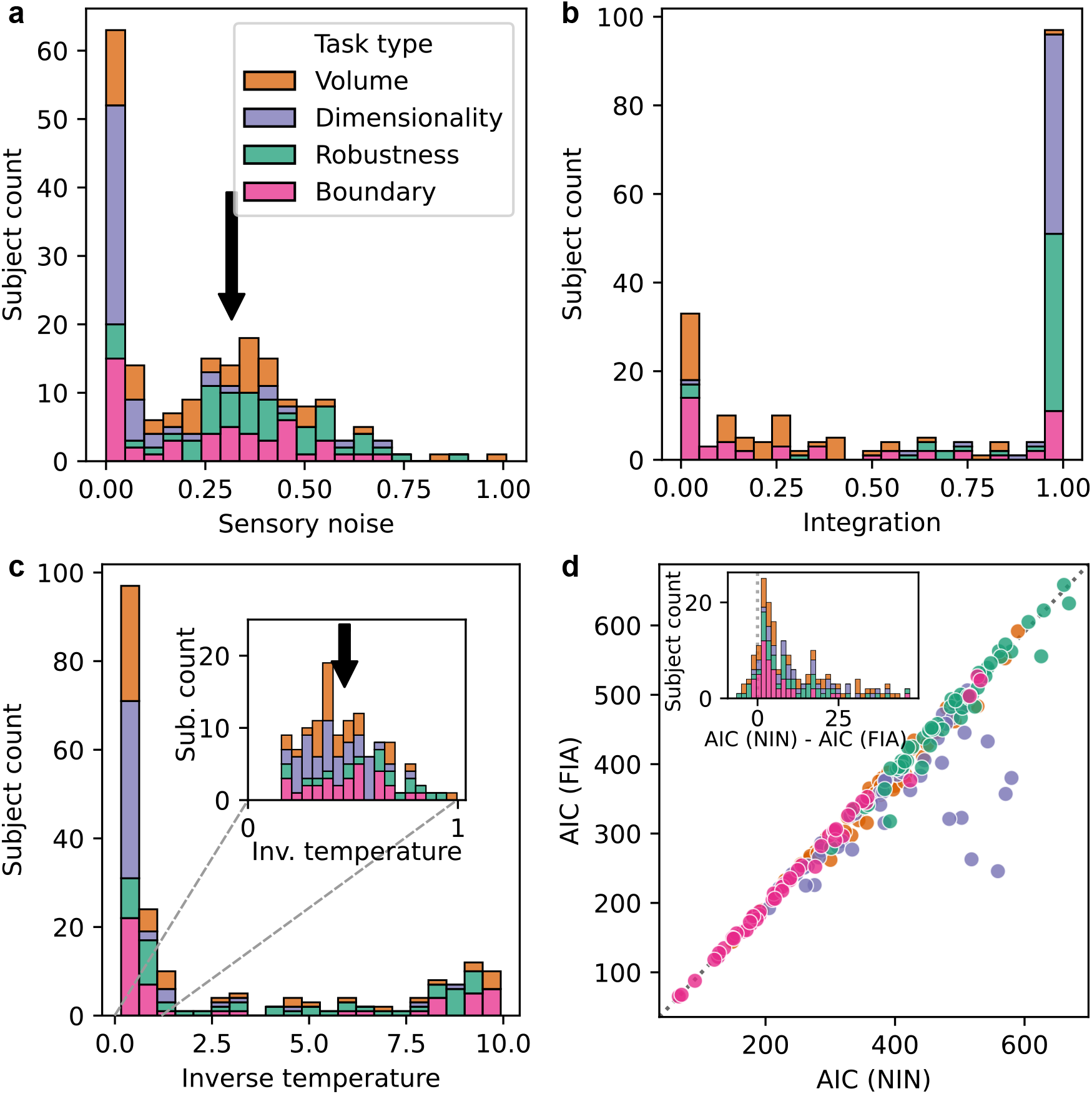
Analysis of human behavior on the generative task, using the Noise-Integration-Noise (NIN) model. **a:** sensory noise (ρ) estimate for all participants, broken down by task type (colors). Arrow: standard error of the location of the centroid of the dot cloud that, on any trial, represented the data X shown to the participants (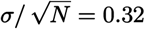, using the notation of A.3 and A.4.2). **b:** estimates of integration parameter b. **c:** estimates of inverse temperature of the choice noise β = 1/T. Inset: detail of the inverse temperature histogram for β ∈ [0, 1]. Arrow: numerical value of the population estimate of likelihood sensitivity from the FIA model. **d:** simple participant-level model comparison (Akaike Information Criterion) between the NIN model and the behavioral model based on the FIA (Equation A106). Lower is better; the dashed diagonal line is the identity. Inset: histogram of NIN-FIA difference, excluding outliers with large positive values, which are overwhelmingly better described by FIA. The AIC is lower (better) for the FIA than for the NIN model for 182 out of 201 subjects.

**Figure B.12.**
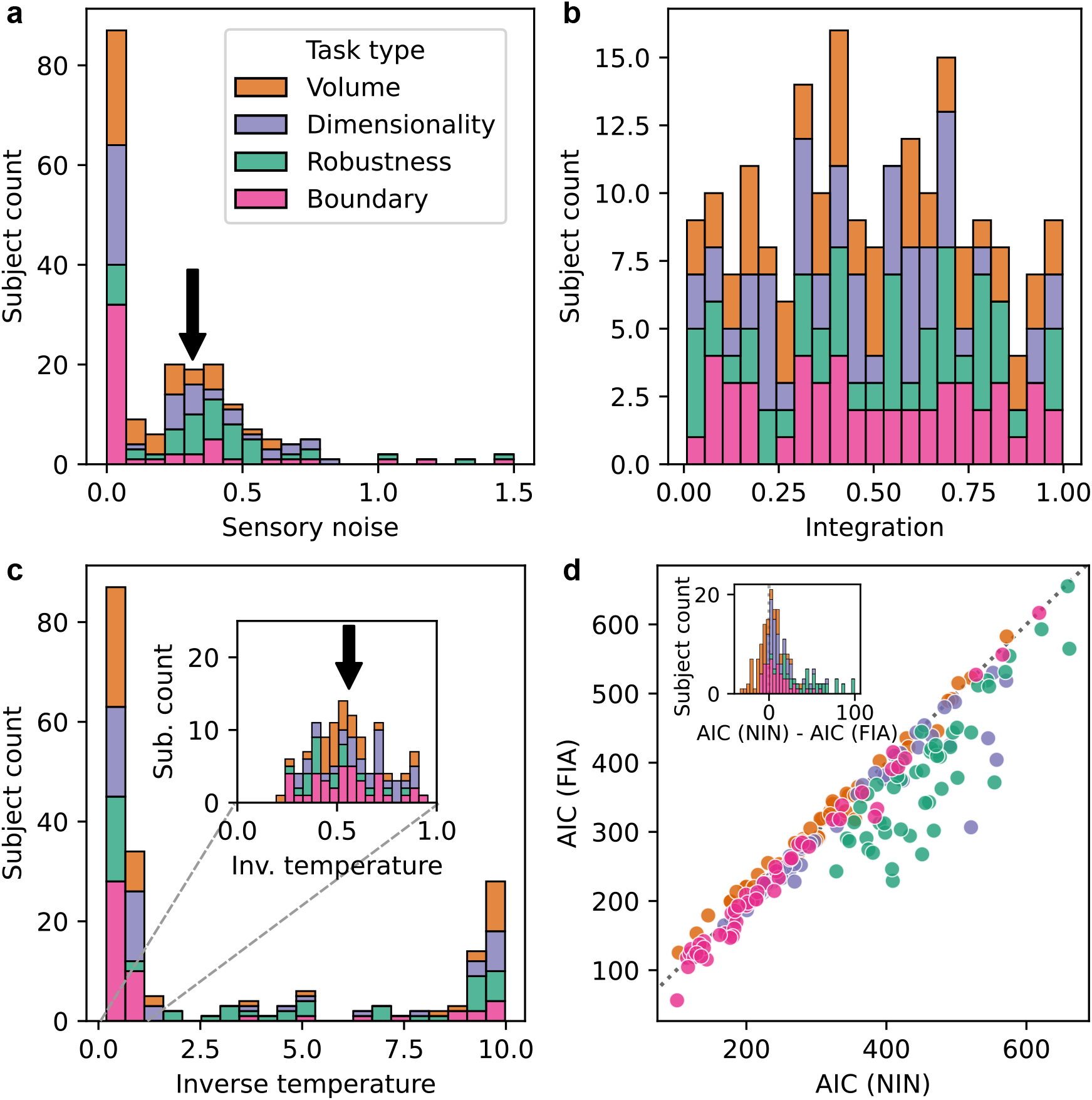
Analysis of human behavior on the maximum-likelihood task, using the Noise-Integration-Noise (NIN) model. Same as Figure B.11, but for the behavioral data of the subjects that performed the maximum-likelihood task. In panel d, the AIC is lower (better) for the FIA than for the NIN model for 144 out of 201 subjects.

